# Phyllotaxis as a dynamical system

**DOI:** 10.1101/2023.02.13.528401

**Authors:** Walch Jean-Paul

## Abstract

One of the major puzzles in phyllotaxis is the much higher frequency of Fibonacci spirals compared to other spiral arrangements such as Lucas spirals. We show that spirals are a form of symmetry, in the same way that axial symmetry is a form of symmetry, which explains why they can be the consequence of many different microscopic phenomena. We apply dynamical systems theory to the main types of phyllotaxis. We show that only Fibonacci spirals should exist and that the other spiral modes (including Lucas) are the consequence of developmental errors, such as the dislocation of a pseudo-orthostichy.

## Introduction

Studying phyllotaxis – its causes, its establishment during development, and its functions – has been a long-standing challenge for scientists and brilliant minds. Historically, Theophrastus, Pliny the Elder, Leonardo Da Vinci, Johannes Kepler and Johann Wolfgang von Goethe were among the most influential pioneering figures in the study of phyllotaxis and paved the way for modern studies. Throughout the 19^th^ and 20^th^ centuries, many hypotheses were tested and numerous articles were published to try to advance our knowledge of the mechanisms underlying different phyllotactic patterns (Florian Jabbour, preface to *Phyllotaxis models*, Walch and Blaise, 2023).

Leaf primordia arise near the apex of the shoot apical meristem (SAM) and move outward and down the dome-shaped meristem. On a cross section of a 15-day old stem apex of *Linum usitatissumum* we can see five clockwise and three anti-clockwise spirals called parastichies (Fig. 1). The number of parastichies is written (5, 3) and is the “mode” of phyllotaxis.

**Figure 1.**
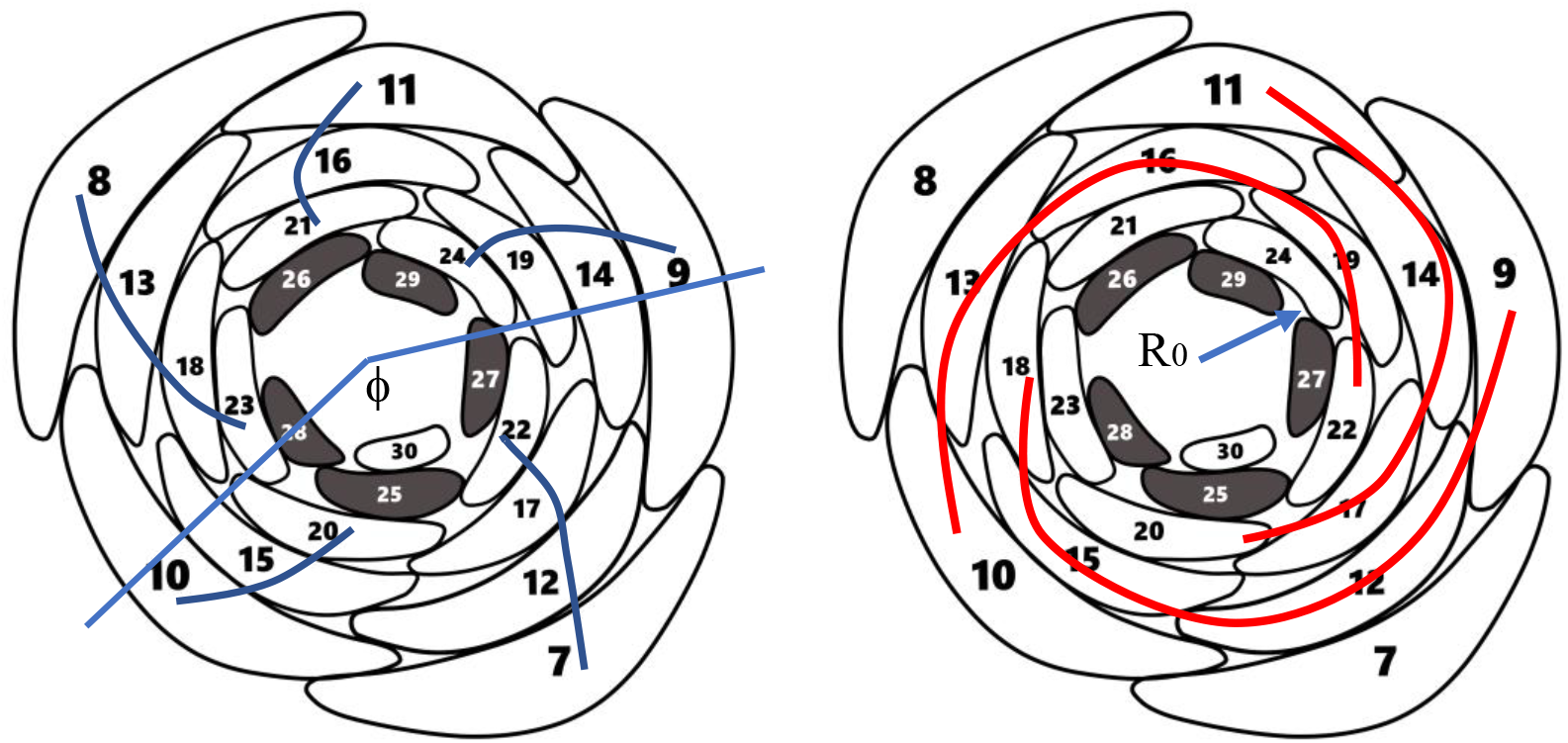
Cross section (centric representation) of a 15-day old shoot apical meristem of *Linum usitatissimum* (from Williams 1975). Numbers correspond to the chronological order of leaf primordia initiation. Parastichies (five clockwise in blue and three anti-clockwise in red) are drawn. ϕ is the divergence angle between the successive primordia (here 9 and 10). Primordia appear at regular intervals on a crown or front (in black: numbers 25-29, primordium 30 is the youngest). R_0_ is the radius of the central zone.

To determine the number of parastichies in a spiral structure, we can either count them or subtract the number of two consecutive primordia on the same parastichy (e.g. 14 – 9 = 5). As the plant grows, the number of parastichies goes from 2, 3, 5, 8, … where each number is the sum of the two previous numbers: this number sequence is the Fibonacci sequence where the ratios of consecutive terms, such as 3/2, 5/3, 8/5,.get closer to the golden ratio (~1.618).

The angle between two chronologically consecutive primordia is called the divergence angle (Fig. 1). In the same way that ratios of Fibonacci numbers converge toward the golden ratio, divergence angles in meristems with numbers of parastichies that are terms of the Fibonacci sequence are close to the golden angle, which is equal to:

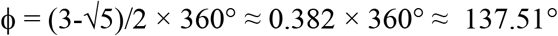

The growth of the SAM can be characterized by the radius R_0_ of the central zone (or of the front), the centrifugal velocity V of the lateral organs moving away from the center as the plant grows, and the time T between the appearance of two consecutive primordia (the plastochron). VT is the distance travelled by a primordium away from the central zone during one plastochron. The ratio of this distance to the radius of the central zone R_0_ gives the growth index G = VT/R_0_, a dimensionless number that characterizes in a unique manner the growth of the SAM, irrespective of size and time scale (Godin et al. 2020). For instance, at day 15, the growth index of *Linum usitatissimum* is 0.07 (Walch and Blaise, 2023).

As the growth rate of the meristem, measured by growth index, decreases with the age of the shoot, the number of parastichies increases. Thus in *L. usitatissimum*, at day 30, the growth index has decreased to 0.04 and the phyllotaxis mode has increased from (3, 5) to (5, 8).

The van Iterson model (van Iterson 1907; Erickson 1973; Godin, Golé and Douady 2020) consists in stacking disks of equal diameter in a cylinder (Fig. 2). From this cylindrical representation, we can observe parastichies (straight lines) that correspond to Fibonacci spirals.

**Figure 2.**
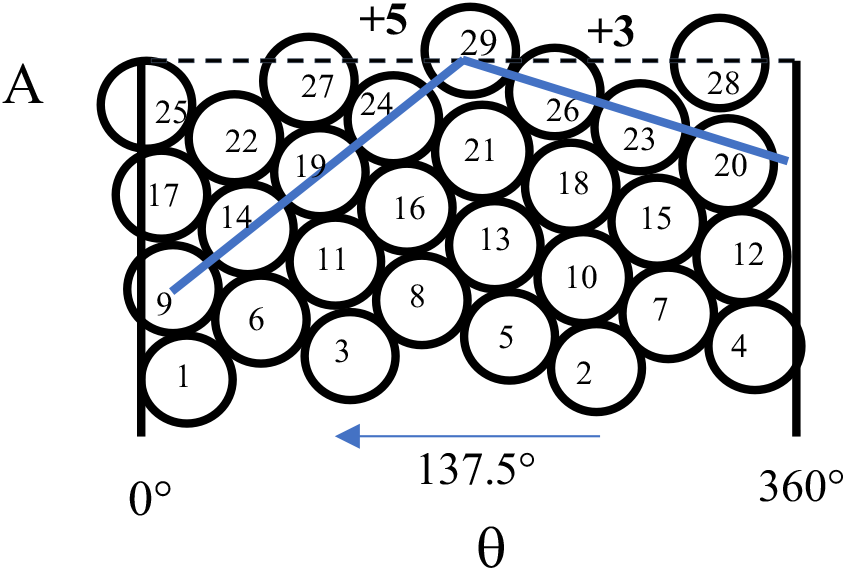
(A) Packing a cylinder with disks of equal diameter: the parastichies (straight blue lines) are here in a (3, 5) mode.

According to Douady and Couder (1992, 1996), an inhibition field produced by young primordia prevents the formation of new primordia for as long as the intensity of the inhibition potential is high enough.

Here, we follow the Douady and Couder model (1992, 1996) and describe the initiation of primordia as depending on the intensity of an inhibition potential at the meristem front. The meristem front is represented by a circle of fixed radius R_0_ = 1 chosen as the unit of length, and primordia are points located between this front and the outer edge of the meristem. The initiation of a new primordium (i) takes place on the front at the point of polar coordinates (R_0_, θ_i_), where θ_i_ is the initiation angle that minimizes the inhibition potential Ui generated by all pre-existing primordia. This potential decreases exponentially with the distance from the primordium generating the inhibition field, the decay length being γ.

If d_ij_ is the distance between the new primordium i and any pre-existing primordium j (between 1 and i-1) at a distance R_j_ from the center of the meristem and with an angular coordinate θ_j_:

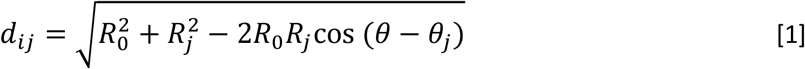

then the potential is given by:

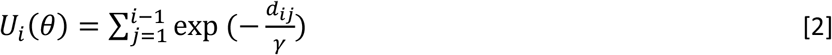

Our model differs from the models of Douady and Couder (1996b), Smith et al. (2006) and Kitazawa and Fujimoto (2015), which all assume that a constant threshold potential value determines primordia initiation and plastochron ratios. Here the model parameters are not the threshold potential value but the plastochron ratios (r) or the distance from the center (see appendix 1):

Since organs are in contact, high plastochron ratios are equivalent to organs being large (Fig. 3A) whereas small platochron ratios (closer to 1) are equivalent to organs being small (Fig. 3B).

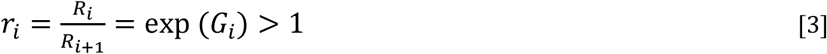

**Figure 3.**
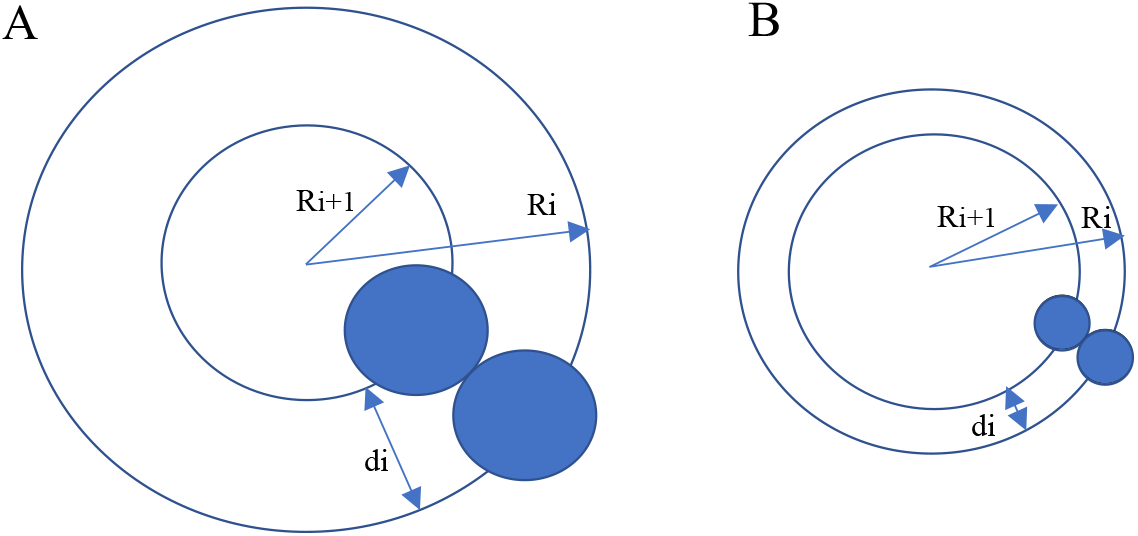
The relationship between the plastochron ratio (R_i_/R_i+1_) and organ size (d_i_). (A) Large plastochron ratio. (B) Small plastochron ratio.

Our model [equations 1-3] allows us to calculate the position of all primordia for given values of the divergence angle and the growth index (Fig.4).

**Figure 4.**
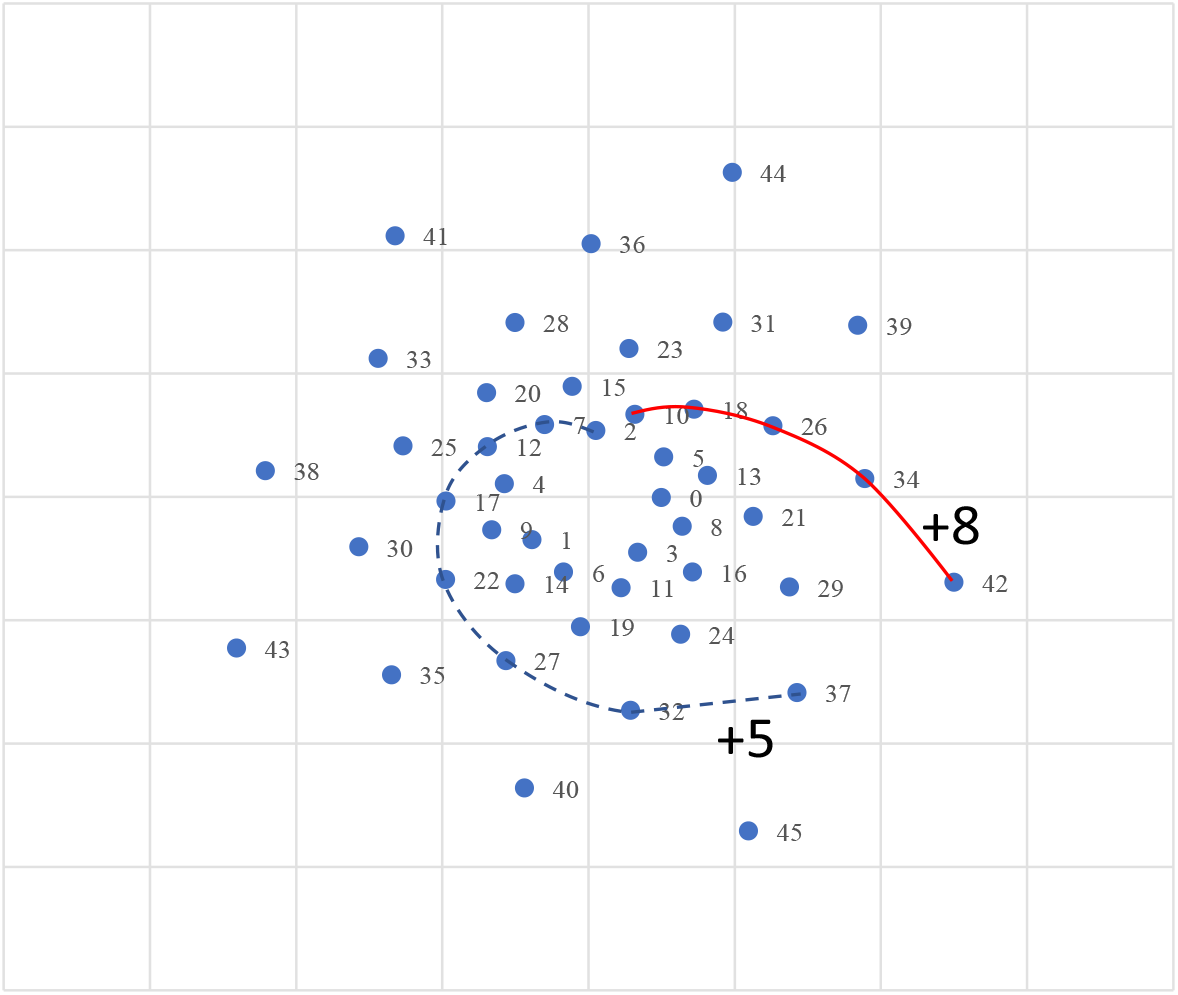
Centric lattice of primordia according to the model for ϕ = 0.382 (the golden angle) and G = 0.04 (corresponding to the (5, 8) phyllotaxis mode). Spirals are +5 and +8 parastichies.

In this paper, we show that spirals are a form of symmetry that is similar to axial symmetry. We apply the mathematical theory of dynamical systems to the main types of phyllotaxis. We show that only Fibonacci spirals should exist and that other types of spirals (including Lucas spirals) are the consequence of errors during development, such as the dislocation of pseudo-orthostichies.

The paper is structured as follows. In section 1, we demonstrate that phyllotaxic spirals are mathematical symmetries. In section 2, we give examples of spirals from the plant world but also from inanimate matter. In section 3, we briefly present Levitov’s energy theory. In section 4, we apply dynamical systems theory to phyllotaxis. In section 5, we develop a stochastic model of phyllotaxis. The discussion focuses on the frequency of Fibonacci phyllotaxis compared to Lucas phyllotaxis.

## 1 Phyllotaxic spirals are a form of symmetry

We define a steady state such that the divergence angle (ϕ) and the growth index (G) are constant. In a cylindrical representation (a Bravais lattice), the points in space (h, θ) are therefore aligned. The line segments thus formed are the parastichies (Fig. 5).

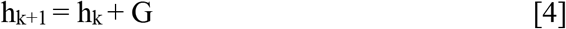

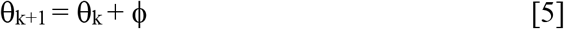

**Figure 5.**
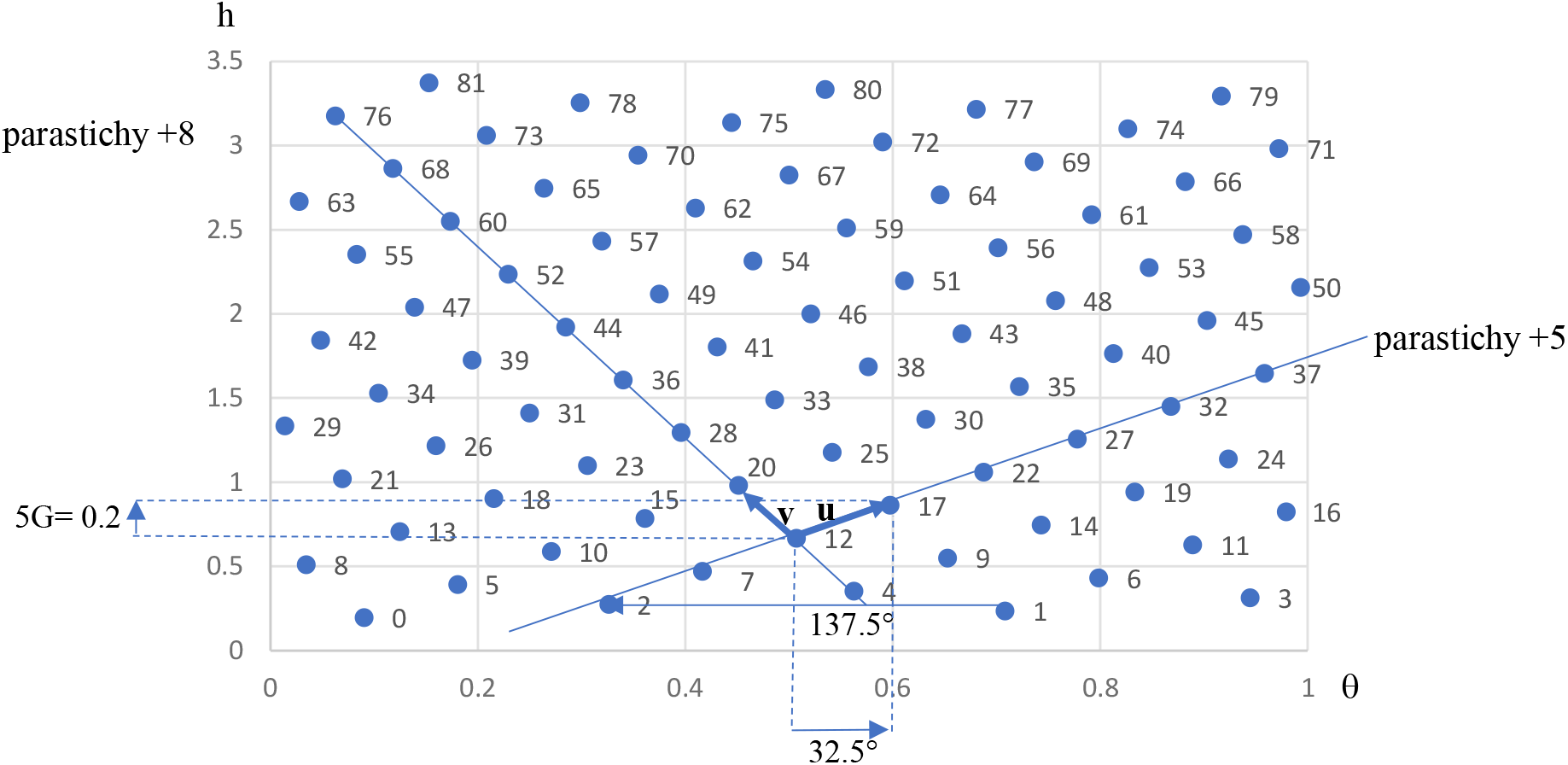
Cylindrical lattice representing the position of 82 primordia for ϕ = 0.382 (the golden angle) and G = 0.04 (phyllotaxis mode (5, 8)).

The correspondence between cylindrical and centric representations was established by Rothen and Koch (1989) (see appendix 2).

A system is symmetric if a transformation applied to all points of the system displaces the intitial points to a place occupied by other points of the system. Thus the corolla of *Lamium galeobdolon* presents axial symmetry, its right side being transformed into the left side (Fig. 6): we say that the corolla of *L. galeobdolon* is invariant by axial symmetry.

**Figure 6.**
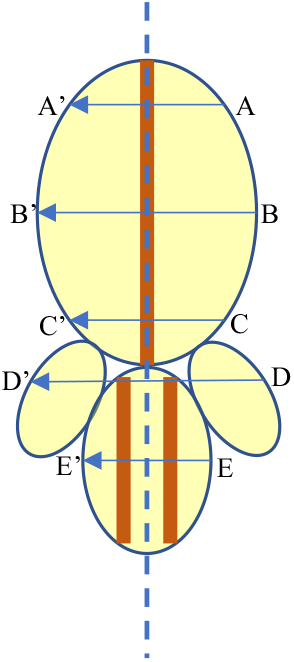
Axial symmetry of the corolla of *Lamium galeobdolon.*

A translation by vector **u** or **v** leaves the cylindrical lattice in Figure 5 invariant: all points of the transformed lattice are superimposed on points of the original lattice (e.g. 12 −> 17).

On the centric figure, this translation corresponds to a rotation of −5 x 137.5° = −327.5° = 32.5° and a dilatation of ratio 1.04^5^ (appendix 2). Thus, point 12 in Figure 7 is in the same place as point 17 in Figure 4.

**Figure 7.**
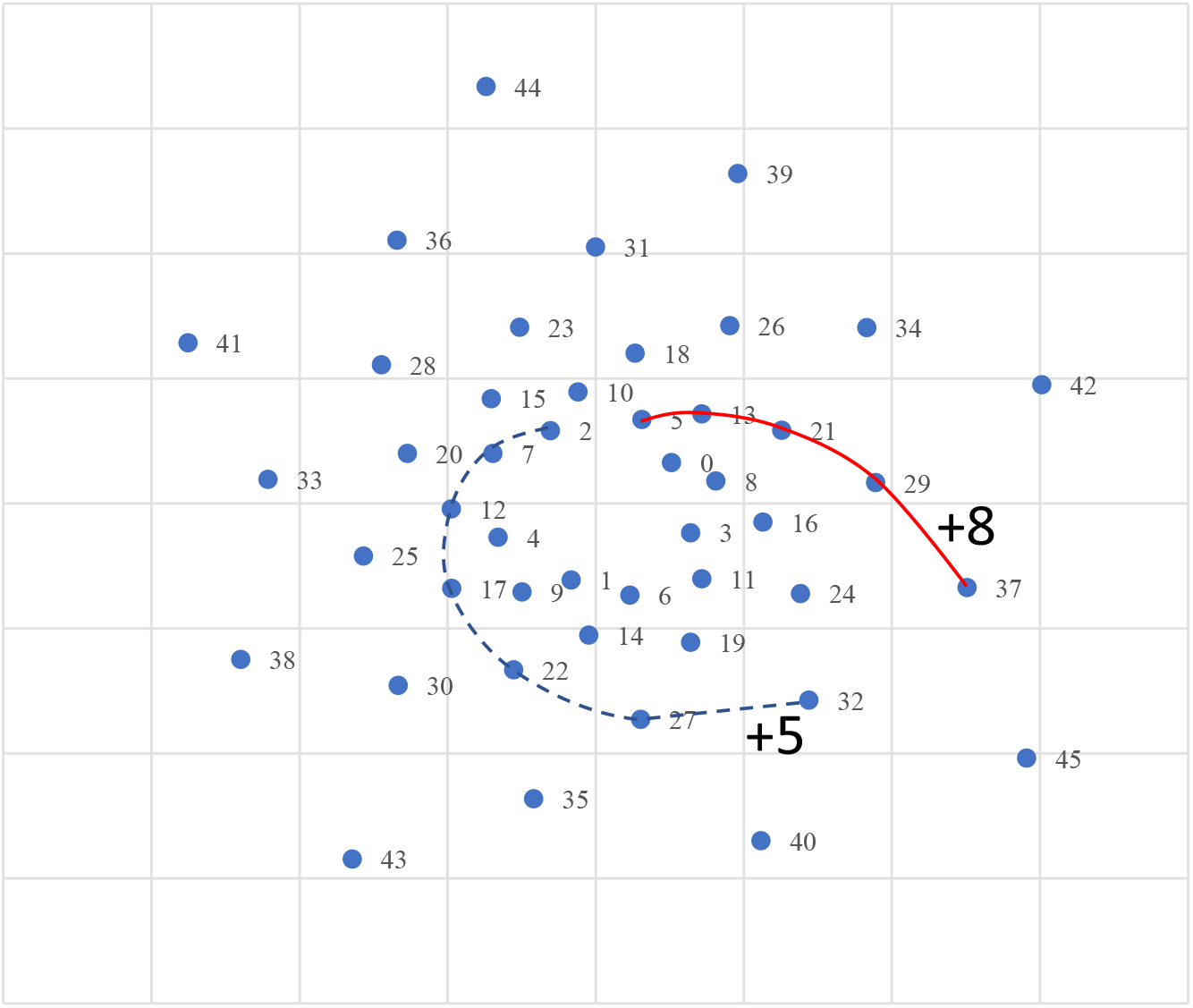
Transformation of the centric representation in Figure 4 by a rotation of 32.5° and a dilation of ratio 1.04^5^.

Thus, for a constant divergence angle and growth index, phyllotaxis is invariant under two transformations corresponding to the two phyllotaxis modes (e.g. +5 and +8 on clockwise and anti-clockwise parastichies). These two transformations, and only these two, transform the two sets of parastichies into themselves.

## 2 Examples of spiral symmetry

### 2.1 Plant lattices

*Lepidodendron* (from the Greek λεπíς scale, and δέvδρov, tree) is an extinct genus of primitive vascular plants known as scale trees or arborescent lycophytes. They thrived during the Carboniferous Period (360 to 300 million years ago) and are part of the coal forest flora. They grew up to 50 meters in height and 1 m in diameter. Fossilized scales at the base of their leaves on the stem show spiral patterns (Fig. 8).

**Figure 8.**
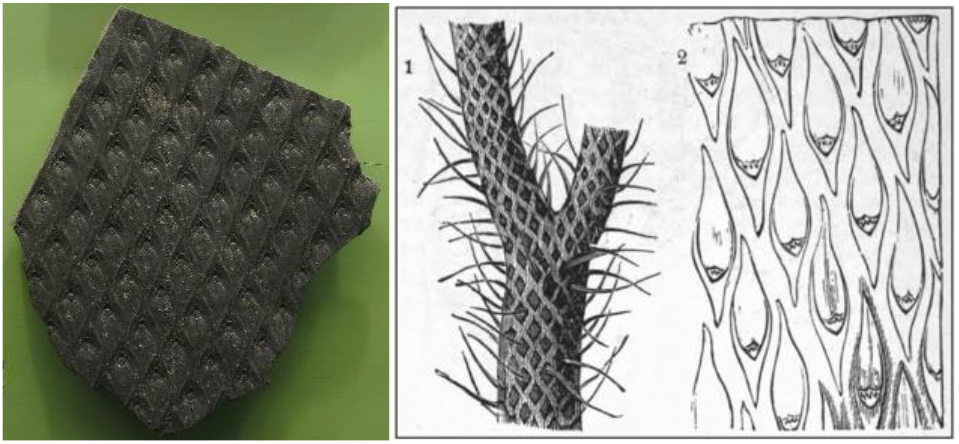
(A) A *Lepidodendron* fossil on display at the State Museum of Pennsylvania. (B) drawing of a Lepidodendron stem from The American Cyclopedia (1879).

The fossil remains of *Lepidodendron* from coalfields in the Glasgow area were studied by Dickson (1871). He determined leaf phyllotaxis in 13 specimens with more or less cylindrical stems. Three of these presented Fibonacci spirals (i.e. (8, 13), (13, 21), (55, 89)), four had multijugate Fibonacci phyllotaxis (*e.g.* (15, 24)), one showed Lucas phyllotaxis (18, 29) and the rest had (9, 14), (12, 19), (10, 13) and (23, 35) phyllotaxis modes.

According to Dickson, the phyllotaxis of *Lepidodendron* is more variable than that of living plants such as *Cactus.*

In angiosperms, the phyllotaxis of leaves (Fig. 1), and subsequently of flowers (Fig. 9), is established in the apical meristems. In the case of *Magnolia soulangeana*, phyllotaxis of the flower is the same as that of the stem (Fig. 10).

**Figure 9.**
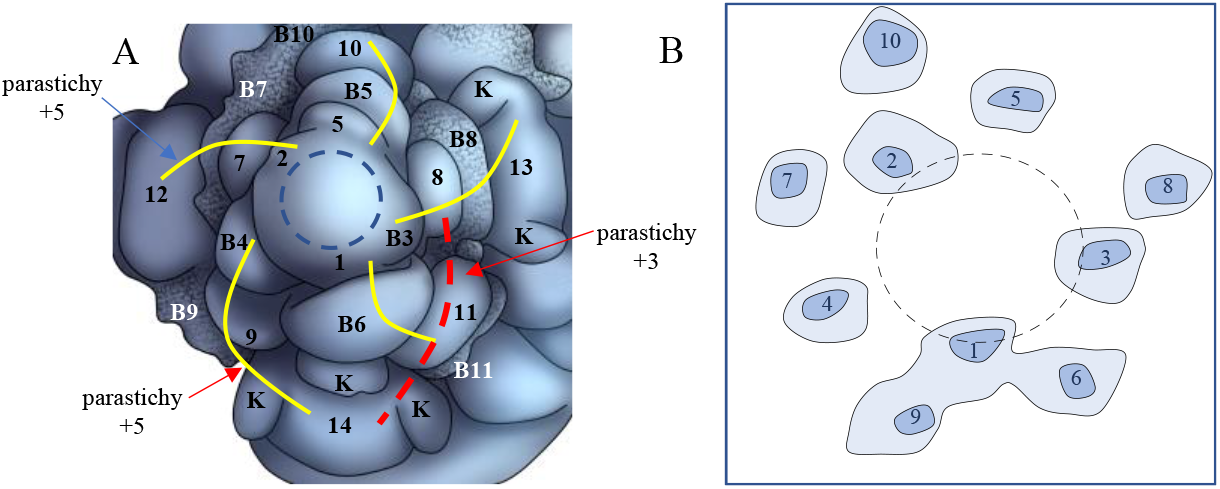
(A) SEM of inflorescence apex of *Antirrhinum majus:* numbers correspond to the nodes, 1 being the youngest; B_n_ corresponds to the bract number; bracts B_7_-B_10_ were removed to make the floral meristems visible; K = sepals. In yellow the five anti-clockwise parastichies; in red, one of the three clockwise parastichies. Redrawn from Carpenter et al. (1995). (B) Model of the *A. majus* raceme (Walch and Blaise, 2023).

**Figure 10.**
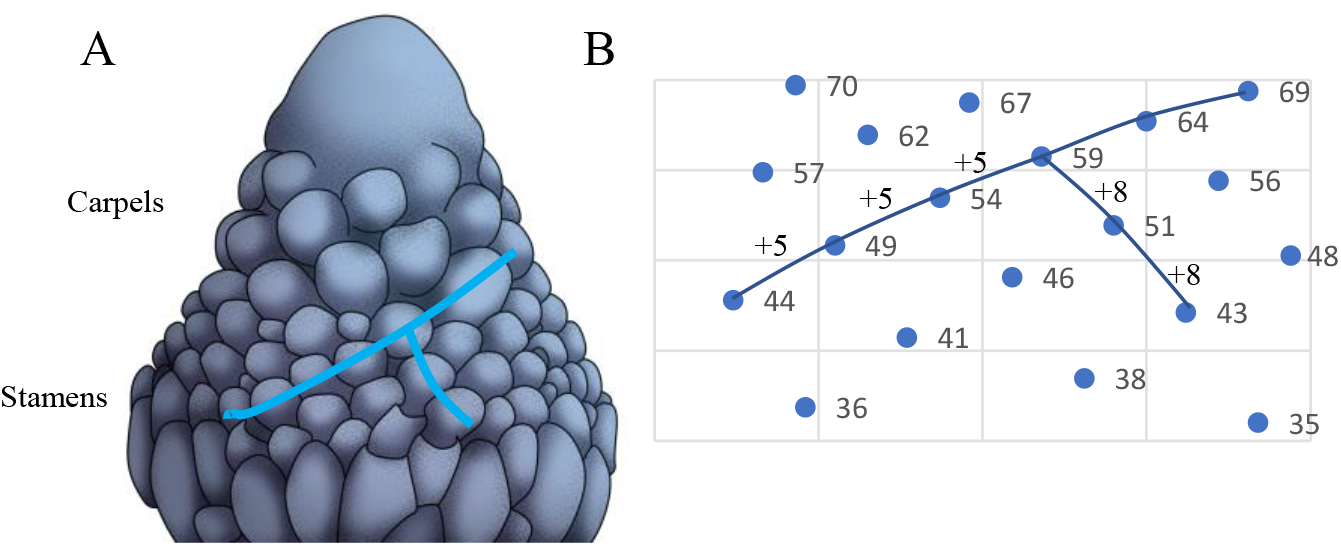
(A) The floral apex of *Magnolia soulangeana* during stamen and carpel initiation, redrawn from Zagórska-Marek (1994). Two partial parastichies are shown. (B) Model of a cylindrical representation of floral organ initiation in *M. soulangeana* (Walch and Blaise, 2023). The same parastichies as in A are shown.

The inflorescence of snapdragon, *Antirrhinum majus* (Plantaginaceae), is a simple raceme with an apical meristem that produces floral meristems following a (3, 5) Fibonacci phyllotaxis.

The floral apex of *Magnolia soulangeana* shows a quite regular Fibonacci pattern with a (5, 8) mode (Zagórska-Marek 1994) (Fig. 10A).

### 2.2 The Douady-Couder experiment

In the Douady-Couder experiment (Douady and Couder, 1996), a horizontal teflon dish filled with silicone oil is placed in a vertical magnetic field (Fig. 11). This field has a weak radial gradient: it is minimal at the center and maximal at the periphery of the dish. Organ primordia are simulated by drops of ferrofluid: in a magnetic field, they are polarized and form dipoles that are attracted toward the region of maximum field. The drops thus spontaneously drift from the center to the border of the dish and repel each other. The repulsion caused by previous drops forces the new drop to place itself in the largest available space at the periphery of the central region.

**Figure 11.**
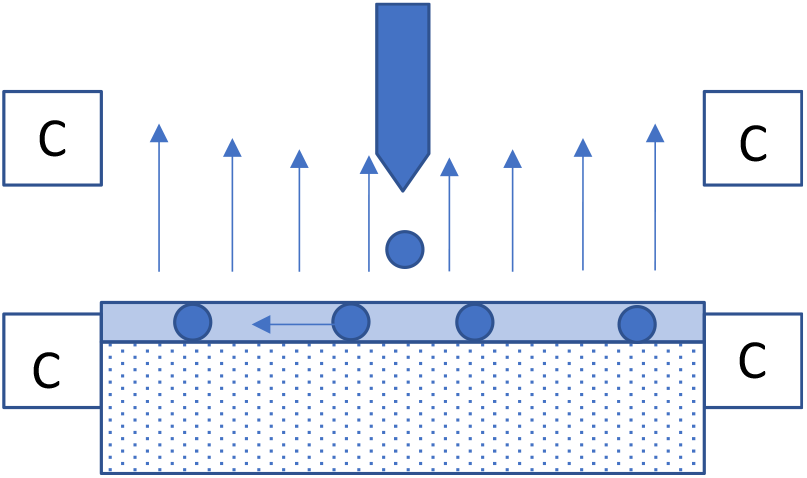
The device used in the Douady-Couder experiment. C = two Helmholtz coils producing a magnetic field. Vertical arrows = magnetic field. Circles = drops of ferrofluid. Horizontal arrow= centrifugal movement of the drops.

**Figure 12.**
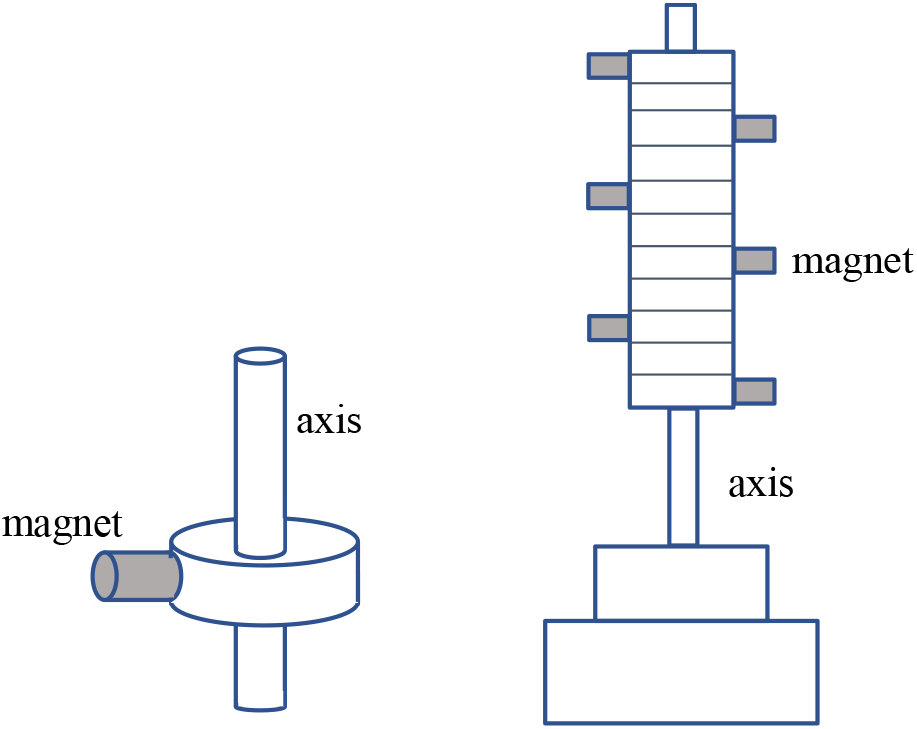
The experimental magnetic cactus apparatus. (A) Each unit of the magnetic cactus consists of a magnet element and a ring secured to the central axis. (B) Schematic representation of the mounted magnetic cactus. From Nisoli et al. (2009).

**Figure 13.**
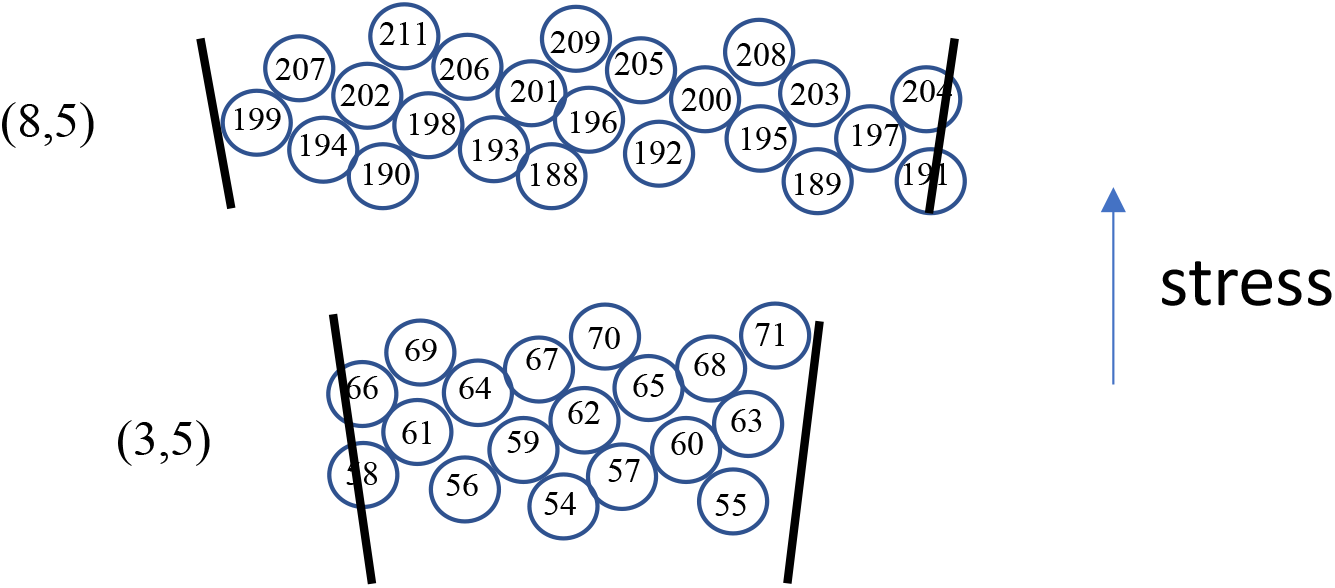
Rising Fibonacci structures (from Atela 2011).

As these drops form identical parallel dipoles, they repel each other with a force that is proportional to d^-4^ (d being the distance between drops). This system gives rise to self-organizing spiral structures. The observed patterns (*e.g.* Fibonacci spirals) depend of the periodicity of drop deposition (T), the radius of the circle outside of which the angular position of the particles does not change (R_0_), and the advection velocity V controlled by the magnetic field gradient.

### 2.3 The magnetic cactus

The “magnetic cactus” consists of equally spaced permanent magnets mounted along an axis around which they are free to rotate (Nisoli et al. 2009). All magnets are outward facing to produce a repulsive interaction between magnet pairs. At the beginning of each experiment the magnets are positioned in a randomly way. They then position themselves and in the final state they give rise to spiral structures with angles of divergence between two successive magnets that are equal to 137.5°, 99.5° or even 77.96° depending on the length of the magnets.

Nisolli and al. used a magnetic dipole-dipole interaction and showed that the lattice energy was minimal for the three so-called (by Jean, 1994) « noble » divergence angles.

## 3 Levitov’s energy model

Levitov modeled a regular lattice of interacting objects that repulse each other (1991, 2021). Lattice geometry is controlled by the balance between external forces and repulsive interactions between different lattice points. The repulsive interaction is determined solely by the distance d between interacting points. It can be, for example, exponential e^-d/γ^ or a power law d^-γ^.

Levitov demonstrated that this energy is invariant under the modular transformation SL(2, Z):

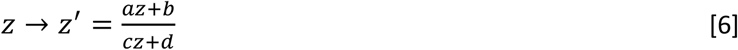

with

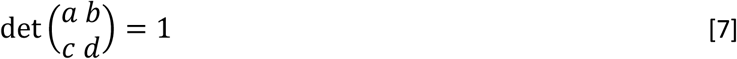

With an increase of the external stress, the lattice structure climbs the Fibonacci sequence.

## 4 Phyllotaxis as a dynamical system

### 4.1 The dynamics of phyllotaxis

We calculated the values of the inhibition potential (U) from the divergence angle (ϕ) and the plastochron ratio (r) using equations [1-3].

In dicots, stem phyllotaxis begins with two opposite facing cotyledons (ϕ = 180°). This state is stable (the potential is minimal) until the plastochron ratio reaches a critical value (r ≈ 1.4) from which two minima arise corresponding to the two opposite chiralities (clockwise and anti-clockwise). Phyllotaxis becomes spiral and its mode is (1, 2) (Fig. 14). This type of bifurcation of a potential is well-known in physics under the name of Hopf bifurcation, which applies to second order phase transitions with a change in the symmetry of the system, e.g. the Rayleigh-Bénard convection. Here, it marks the transition from a verticillate distichous phyllotaxis to a spiral phyllotaxis. The two chiralities are equiprobable.

**Figure 14.**
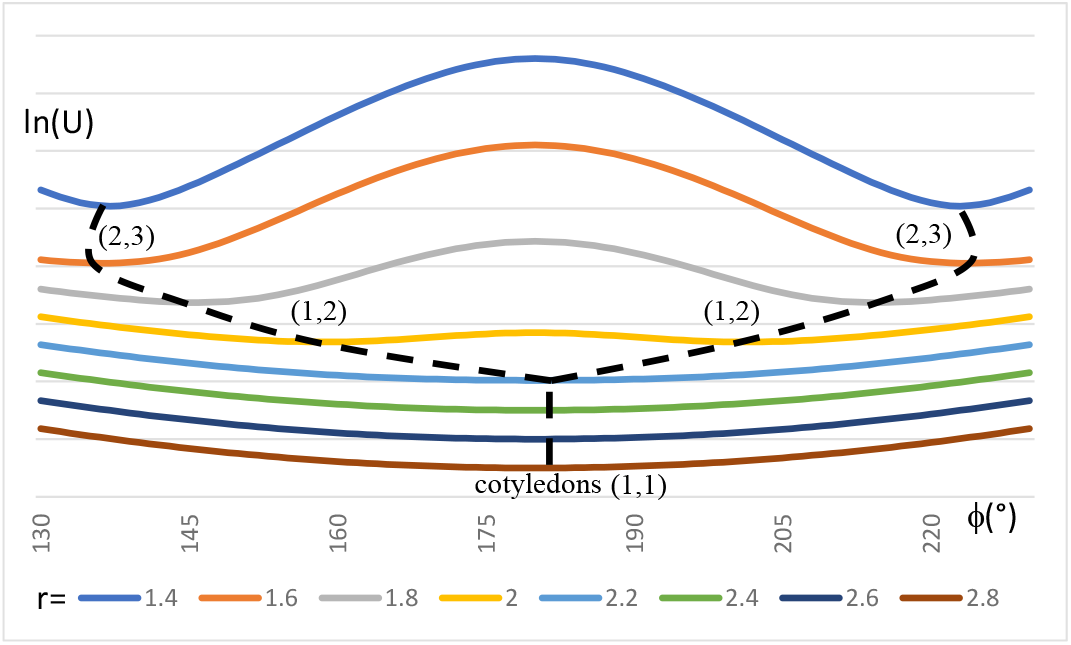
The inhibition potential (ln(U)) as a function of the divergence angle (ϕ) for different plastochron ratios (r) calculated from the model. For each value of r, the curve of the natural logarithm of the inhibition potential is shown; trajectories in phase space as r decreases climb the valleys formed by the inhibition potential curves (dashed lines).

In the vast majority of plants with spiral phyllotaxis, the number of parastichies follows the terms of the Fibonacci sequence. Some plants, however, are arranged according to other sequences, and the most common of these is the Lucas sequence (2, 1, 3, 4, 7, 11, …).

Primordia initiate along the valleys (local minima) of the inhibition potential. The dynamics of the system as the plastochron ratio decreases is consistent with that of van Iterson’s diagram (Douady and Couder, 1996): the same bifurcations are found, the number of inhibition potential minima increasing with a decrease in the value of the plastochron ratio (Fig. 15). The system rises in the Fibonacci sequence by a succession of quasi-bifurcations. Indeed, at each evolution, such as from (3, 5) to (5, 8), the passage to (3, 8) is impossible because there is a potential barrier (a “mountain”) between the two modes.

**Figure 15.**
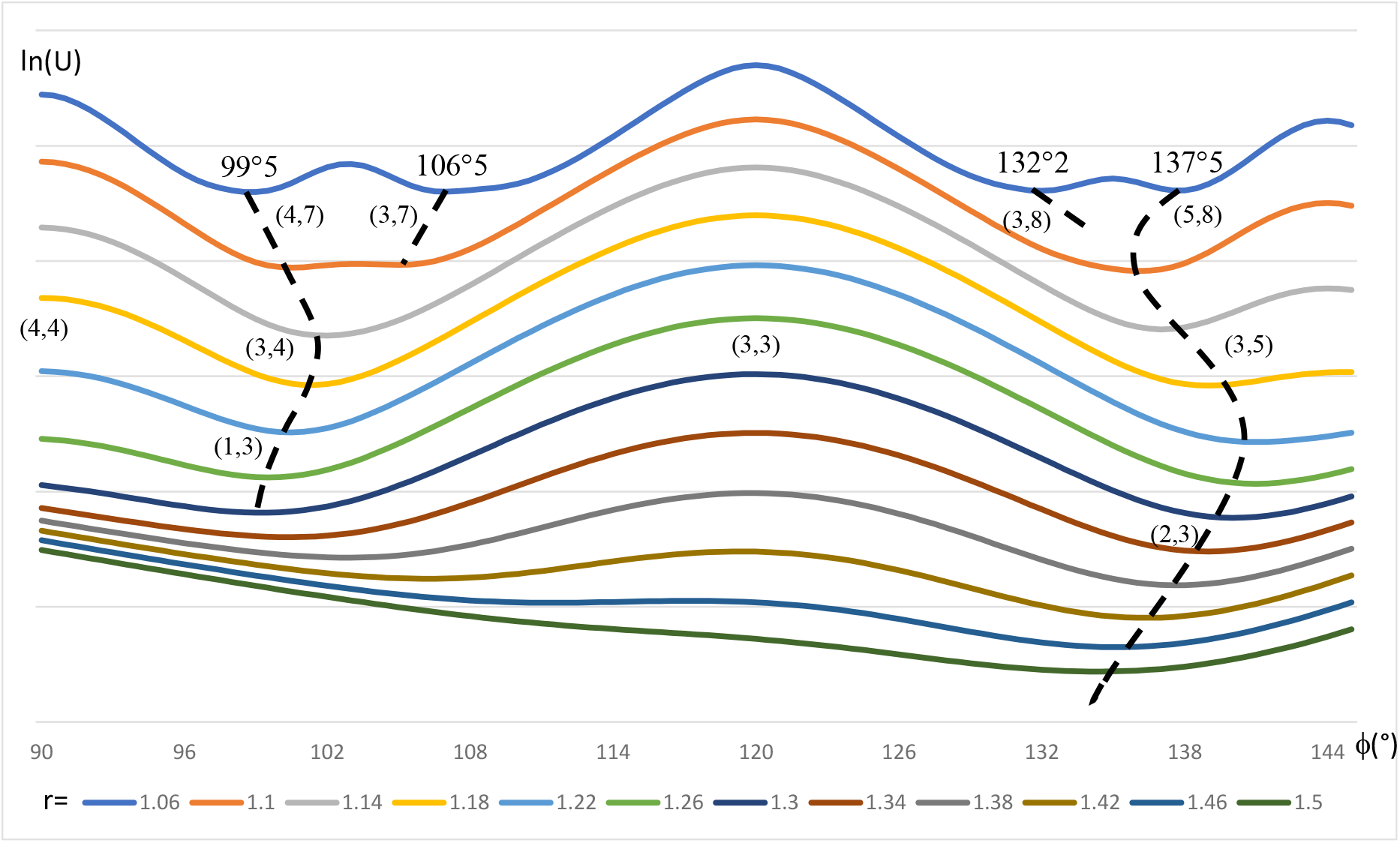
The inhibition potential (ln(U)) calculated from the model as a function of the divergence angle (ϕ) for different plastochron ratios (r). For each value of r, the curve of the natural logarithm of the inhibition potential is shown; trajectories in phase space as r decreases climb the valleys formed by the inhibition potential curves (dashed lines). The angles given above the curve are the noble angles toward which each trajectory converges. (4, 4) and (3, 3) are whorled tetramerous and trimerous phyllotaxis (ϕ= 90° and ϕ= 120°), respectively.

Only Fibonacci phyllotaxis can develop in continuity with the first two cotyledons. Indeed, when plastochron ratios decrease, only quasi-bifurcations appear, the minimum potentials of the branches leading to angles of divergence that are different from the golden angle appearing in discontinuity with the parent branch.

This is the reason for the very high frequency of Fibonacci phyllotaxis compared to other types of phyllotaxis. “Accidents” must happen during development for other types of spirals to occur in a plant, the most frequent being Lucas phyllotaxis, the first to appear (of mode (1, 3)) for a plastochron ratio r ≈ 1.3.

### 4.2 Whorled phyllotaxis

Spiral arrangements are not the only type of phyllotaxis. Whorled patterns consist of alternating whorls, each composed of the same number of organs (merosity). Divergence angles between organs within a whorl are constant and equal to 360° divided by the number of organs: for instance the divergence angle is = 360°/2 = 180° in a dimerous whorl, 360°/3 = 120° in a trimerous whorl and 360°/4 = 90° in a tetramerous whorl. Whorled patterns present perfectly vertical orthostichies on the shoot unlike the inclined pseudo-orthostichies of spiral phyllotaxes.

The vegetative meristem of *Plantago webbii* (Plantaginaceae, Lamiales) is tetramerous. The four members of one whorl are initiated simultaneously at the shoot apex (Fig. 16) (Rutishauser 1998). Each whorl has the same number of primordia.

**Figure 16.**
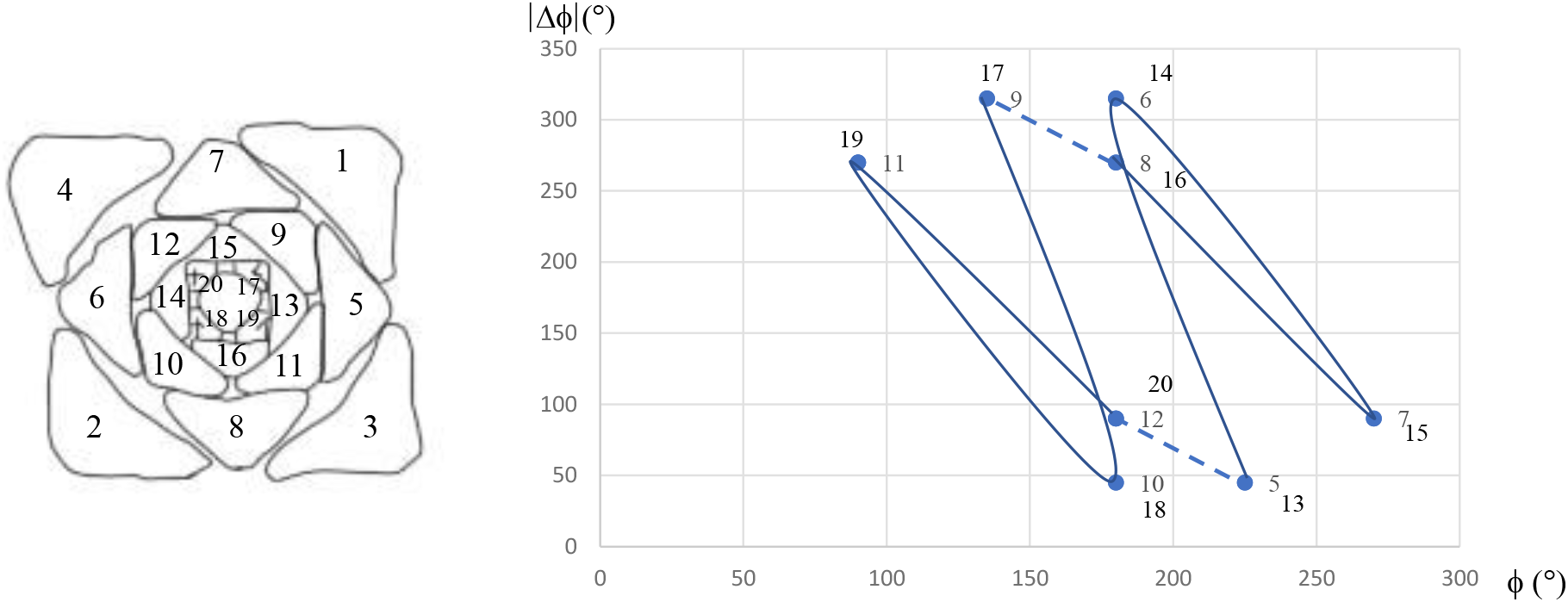
(A) Cross section of the apical vegetative meristem of *Plantago webbii* (Plantaginaceae), from Rutishauser (1998). (B) Phase portrait of *Plantago webbii.*

In dynamical system theory, the phase portait represents the evolution of a system in phase space, here (ϕ, Δϕ) where ϕ is the divergence angle between points i+1 and i and Δϕ is the difference between the divergence angles at point i+1 and point i (we have chosen abs(Δϕ) for more visibility).

The whorled vegetative meristem of *Plantago webbii* walks on a “butterfly” of eight points and comes back to each point at each orthostichy (Fig. 16). In physics, the archetype of a periodic phase portrait is a simple pendulum without friction.

### 4.3 Fibonacci phyllotaxis

The plastochron ratios measured in the vegetative meristem of *L. usitatissimum* (Williams 1975), with Fibonacci phyllotaxis, can be approximated by a decreasing exponential function of organ number (n):

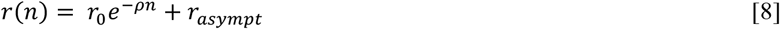

with r_0_ = 0.35 (the initial value), ρ = 0.18 (the decrease rate) and r_*asympt*_ = 1.05 (the level of the asymptote) (Blaise and Walch, 2023, chp. 6). With these parameters and starting with two opposite primordia, we obtained a rapid convergence of the divergence angles toward the golden angle and a (5, 8) phyllotaxis mode. We call these parameters “the canonical parameters”: Fibonacci phyllotaxis requires a “slow and contolled” decrease in plastochron ratios.

We can compare the result of the model (Fig. 17A) with Fig. 1 (redrawn from Williams (1975)) and appreciate how well the model fits the observations of this vegetative meristem.

**Figure 17.**
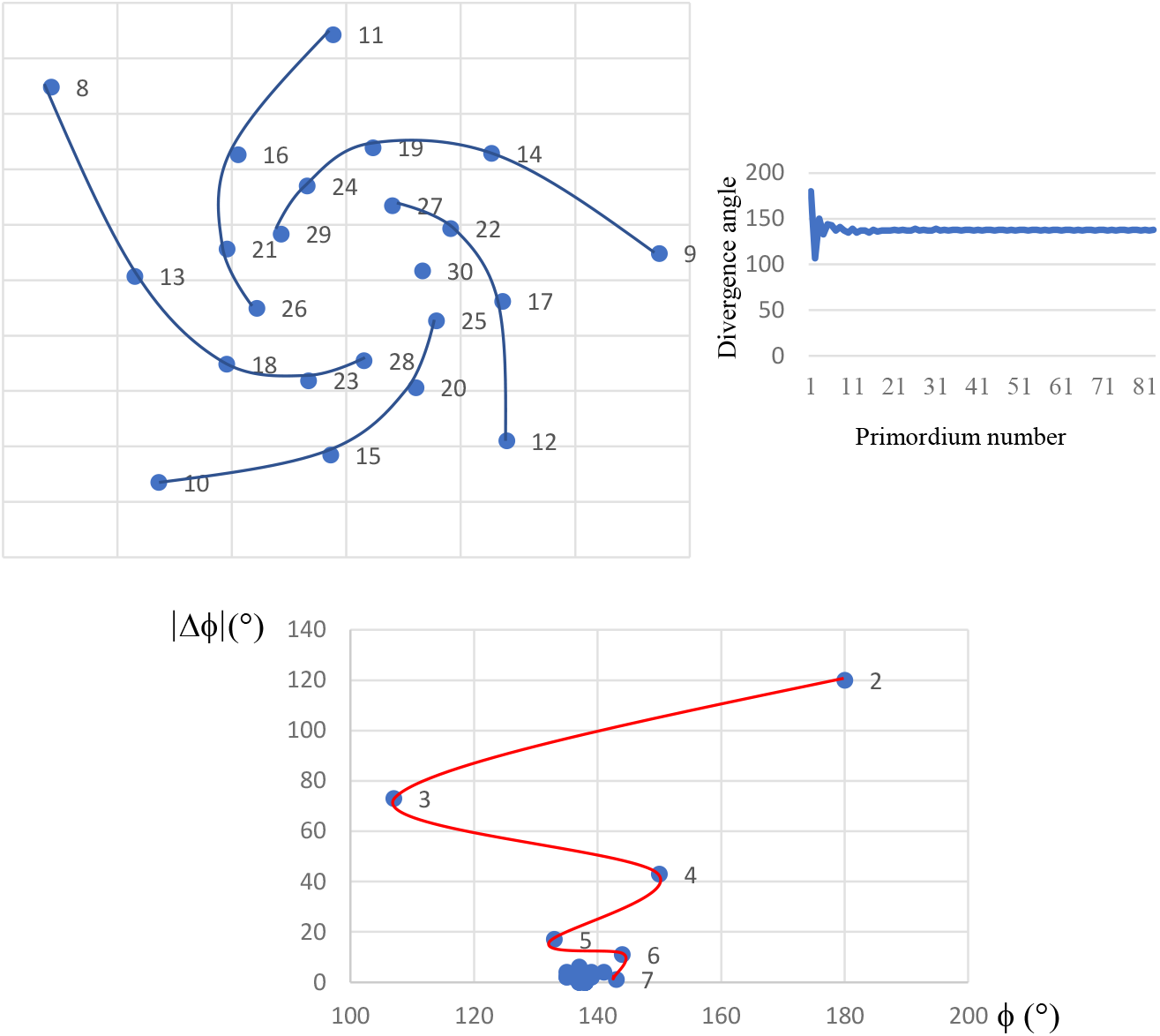
(A) Model of the vegetative meristem of *L. usitatissimum* at day 15 of development using the canonical parameters (r_0_ = 0.35, ρ = 0.18 and r_asympt_ = 1.05). (B) Value of the divergence angle between successive primordia derived from the model with the canonical parameters for 81 primordia. (C) Phase portrait of Fibonacci phyllotaxis. Numbers correspond to the chronological order of leaves initiated along the shoot.

The trajectory of the phase portrait of the Fibonacci vegetative meristem of *L. usitatissimum* converge to the golden angle. In physics the archetype of a convergent phase portrait is that of a pendulum with friction whose oscillations are damped. The golden angle is called a fixed point or attractor toward which the trajectory of the system converges.

### 4.4 Periodic divergence angles

If plastochron ratios decrease faster than the canonical value (ρ = 0.3 vs. ρ = 0.18), we observe a permutation of organs: for instance, the divergence angle between points 31 and 32 is 140° (anti-clockwise), a value that is close to the golden angle, but that between points 33 and 34 is 360° – 86° = 274° (anti-clockwise). Organ number 34 should be located +137° from point 33, however that is where point 35 is located, point 34 jumping instead to the next location and leaving its palce to point 35.

As primordia are permutated, divergence angles oscillate between ≈ 137°, 230° and 274° (Fig. 18B) and there are several attractors in the phase portrait. This type of permutation has been observed in *Doryphora aromatica* (Laurales) (Staedler and Endress 2009; Blaise and Walch 2023).

**Figure 18.**
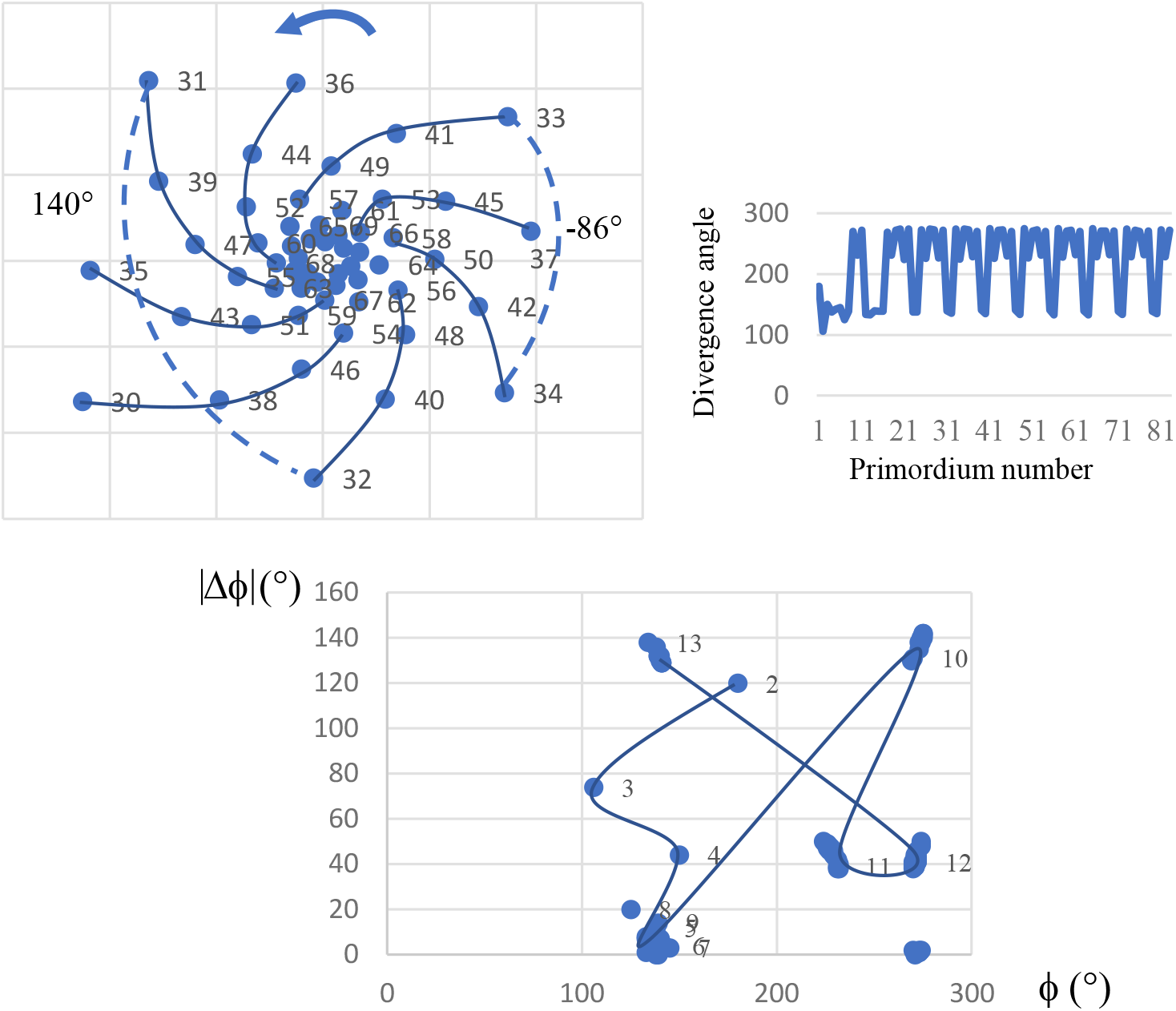
The pattern obtained with r_0_ = 0.35, ρ = 0.3 and r_asympt_ = 1.05: (A) centric representation of organ arrangement; (B) value of the divergence angles showing a periodic pattern; (C) phase portrait of periodic phyllotaxis.

### 4.5 Chaotic/quasi-symmetric phyllotaxis

For r_asympt_ = 1.02 and ρ =0.7, we find nine pseudo-parastichies in both directions (i.e. a (9, 9) mode) (Fig. 19). The inclination of the parastichies in both directions is more or less equal (Fig. 20). Thus phyllotaxis is quasi-symmetric.

**Figure 19.**
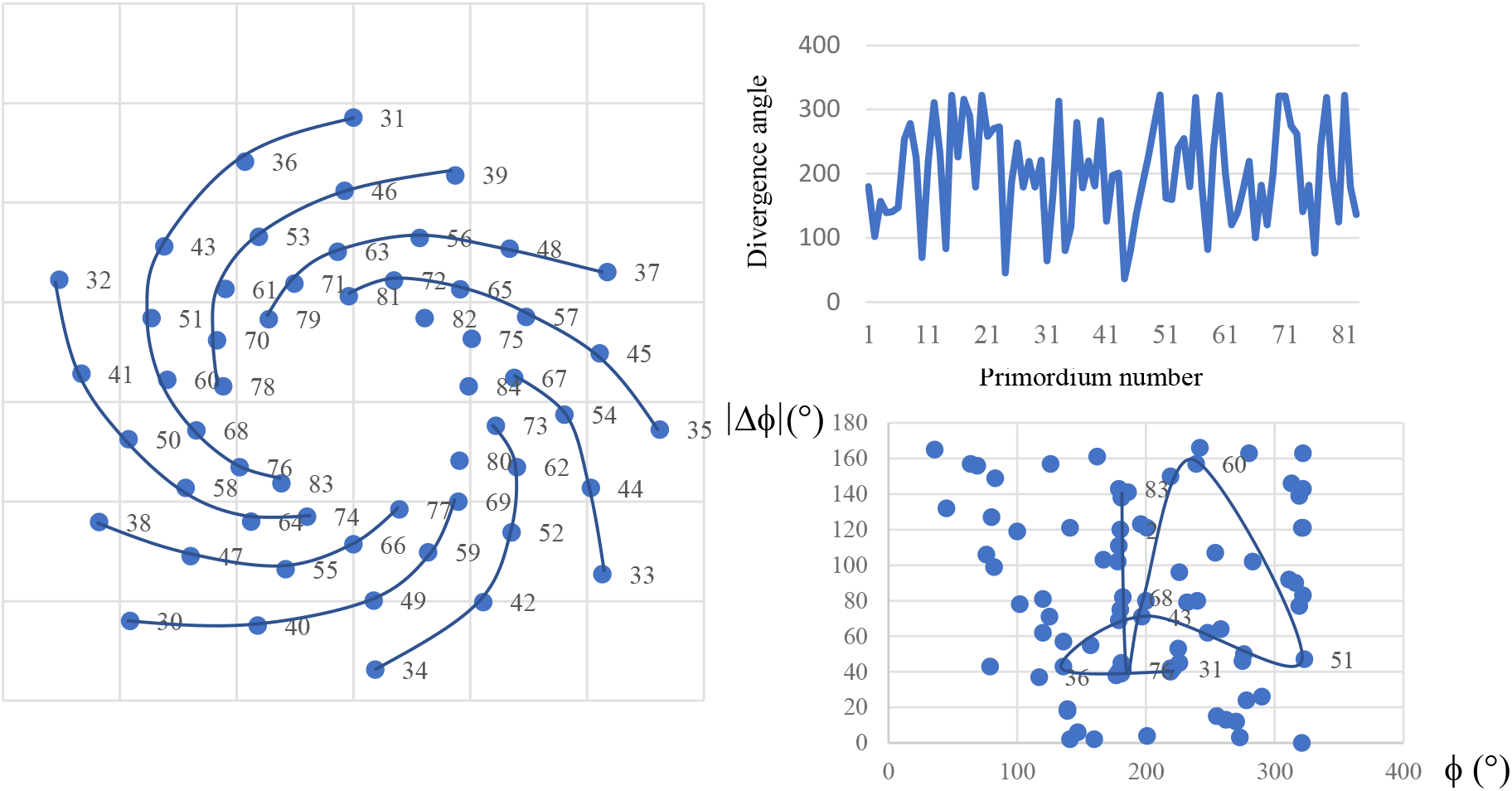
Pattern obtained with r_0_ = 0.35, ρ = 0.7 and r_asympt_ = 1.02: (A) centric representation of organ arrangement; (B) divergence angles showing a chaotic pattern; (C) phase portait of chaotic phyllotaxis.

**Figure 20.**
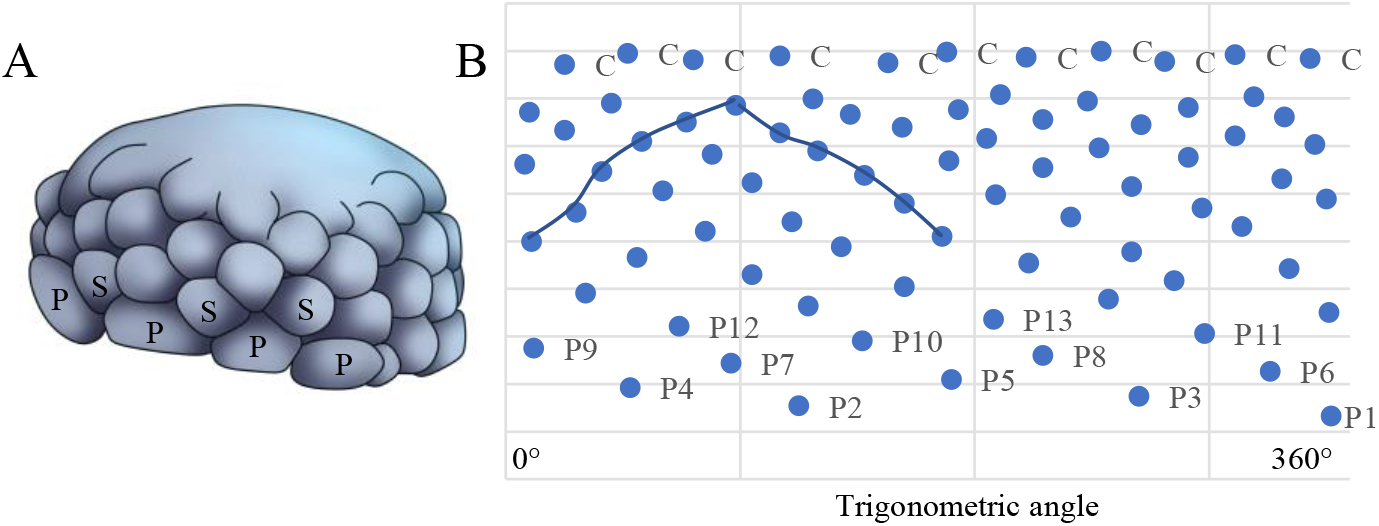
(A) The floral meristem of *Nuphar pumila* (from El et al. 2020); (B) Cylindrical representation of the modeled floral meristem of *Nuphar pumila* showing quasi-symetric pseudo-parastichies (from Walch and Blaise 2022). P = petal (the associated number corresponds to the order of initiation); S= stamen; C = carpel.

Using the disk stacking model, Golé, Dumais and Douady (2016) showed that a rapid decrease in disk diameter led to the breakdown of Fibonacci spirals in favor of a “quasi-symmetric” arrangement, with a nearly equal number of clockwise and anti-clockwise spirals. This arrangement can be seen in inflorescences of *Spathypillium* (Araceae), *Banksia* (Proteaceae), corn (*Zea mais*) and peteh (*Parkia speciosa*, Leguminosae subfamily Mimosoideae) (Godin, Golé and Douady, 2020; Supplementary information).

According to our model, if plastochron ratios decrease very rapidly, phyllotaxis becomes quasi-symmetric with a near equal number of left and right parastichies. This type of spiral phyllotaxis is different from Fibonacci phyllotaxis, and can be mistaken for whorls. Pseudo-parastichies bring out an apparent order in the chaos of divergence angle values.

Floral phyllotaxis in *Nuphar* (Nymphaeaceae) has traditionally been thought to be spiral, a view supported by August Eichler (1878); however, Endress and Doyle (2015) have asserted that all Nymphaeaceae have flowers with complex-whorled (i.e. non-spiral) phyllotaxis. The criterion used by Endress is as follows: phyllotaxis is whorled when the inclination of the left and right parastichies is the same. However, the criterion used by Endress and Doyle is insufficient: an equal inclination of left and right parastichies differentiates whorled phyllotaxis from Fibonacci phyllotaxis, but not whorled phyllotaxis from certain types of irregular spirals.

According to Rutishauser (2016), the cauline phyllotaxis of *Acacia* (Australian “wattles”, Leguminosae subfamily Mimosoideae) can be spiral (Fibonacci system) (*A. amblygona* and *A. rossei*), “complex whorled” (*A. adoxa* with 9 to 10-merous phyllode whorls) or even irregular without any obvious order (*A. sphacelata subsp. verticillata* and *A. conferta)*. The latter are correlated with geometrical parameters such as leaf and stamen primordia that are very small as compared to the size of the apical meristem.

According to Douady and Golé (2016), the male cone of *Cedrus* is a remarkable example of drift from Fibonacci to quasi-symmetric, as is the inflorescence of *Parkia speciosa*, which shows a solid quasi-symmetric pattern originating from a rapid transition.

For r_0_=0.35 and r_asympt_= 1.02, the basin of attraction (the set of initial conditions leading to a long-time behavior that approaches an attractor) of Fibonacci phyllotaxis corresponds to values of the control parameter ρ less than 0.2; that of permuted phyllotaxis to values of ρ between 0.2 and 0.25; that of quasi-symmetrical phyllotaxis to values of ρ greater than 0.25 (Figure 21).

**Figure 21.**
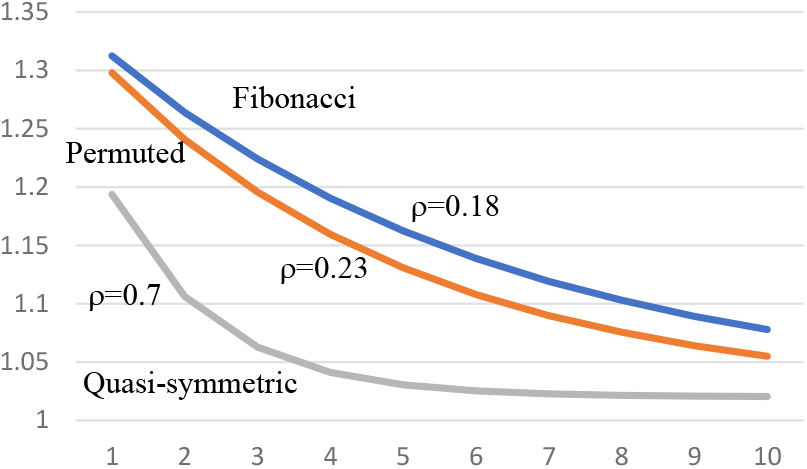
Profile of plastochron ratios for three different decrease rates (r_0_=0.35; r_asympt_= 1.02).

Fluid flow behavior is controlled by similar dynamics. The transition from a laminar to a turbulent flow is determined by Reynolds number, which combines the velocity of the fluid, the width of the river in which it flows, and fluid density and viscosity. When Reynolds number is small, as in a wide, gently sloping river, the flow is laminar. By contrast, a mountain torrent near its source (characterized by fast flowing water and a narrow width) will have a high Reynolds number: the flow is turbulent, and the movement of each drop of water is unpredictable. Between these two regimes, the flow is periodic.

### 4.6 Pseudo-whorled phyllotaxis

Some Magnoliales and Laurales align organ identity with spiral phyllotaxis based on “series” (Staedler and al. 2007) or pseudo-whorls (Walch and Blaise, 2023), i.e. sets of organs of the same type that look like whorls but are less regular because they are initiated in a spiral: the divergence angles are golden angles rather than rational angles that divide a circle into equal parts.

According to Erbar and Leins (1994), the perianth of *Liriodendron tulipifera* (Magnoliales); (K1-K9 in Fig. 22A) shows a transition from spiral phyllotaxis to trimerous whorls. Three short plastochrons are followed by a relatively longer plastochron, which is followed by three short plastochrons, a longer one and again three short plastochrons. Although the perianth shows clear indications of spiral initiation, perianth organs are arranged in trimerous “whorls”. The divergence angles of subsequent primordia (e.g. stamens numbered 1-8 on Fig. 22A) are equal to the golden angle.

**Fig 22.**
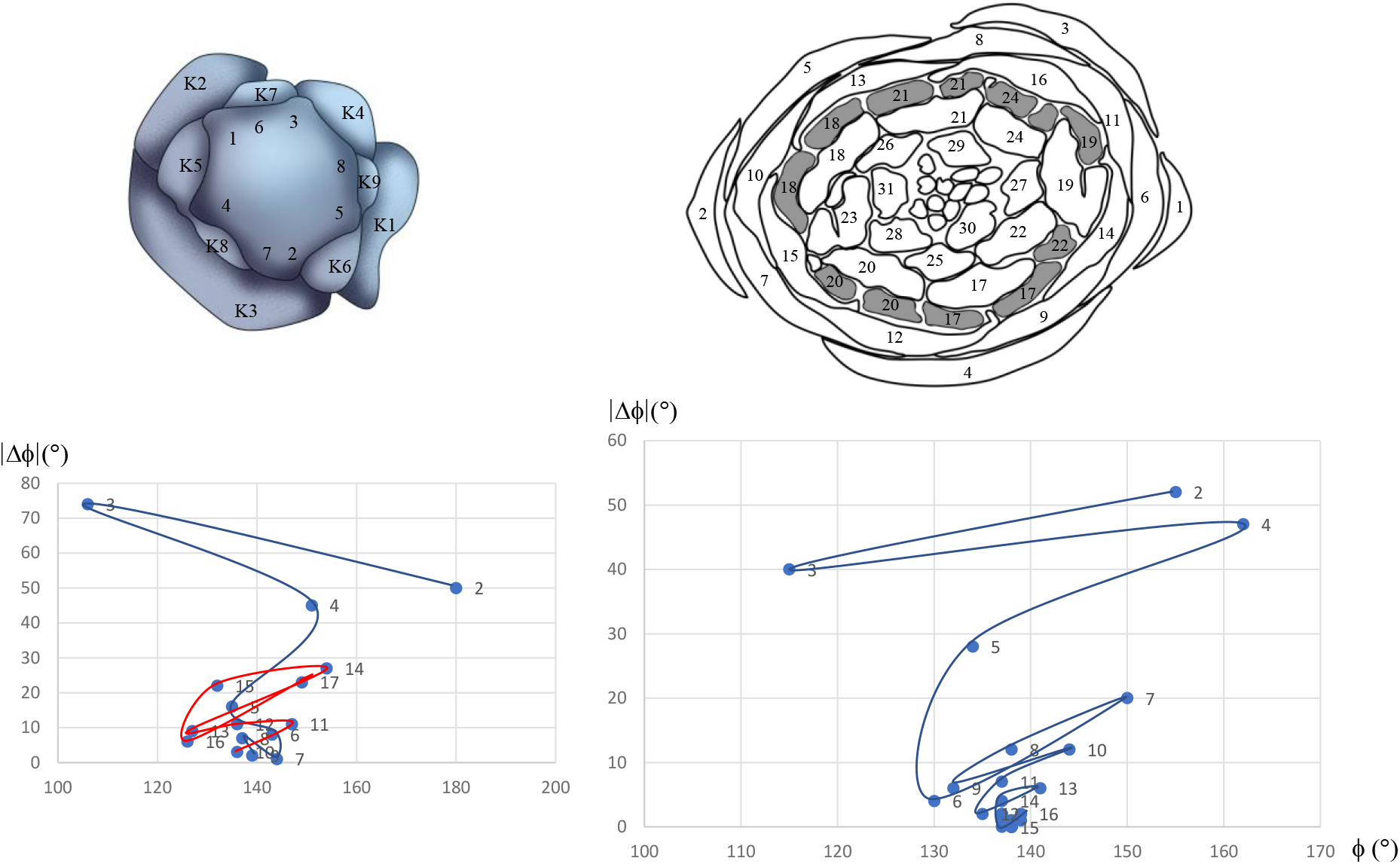
(A) The floral meristem of *Liriodendron tulipifera* redrawn from an SEM micrograph from Erbar and Leins (1994) and (B) phase portrait of floral organ initiation in *L. tulipifera*, with r_0_ = 0.35, ρ = 0.18 and r_asympt_ = 1.02; the plastochron ratio between pseudo-whorls is set to r = 1.30. (C) Transverse section of the flower bud of *Daphnandra repandula*, from Staedler and Endress (2009) showing organs belonging to distinct pseudo-whorls. (D) Phase portrait of *D. repandula* (R_0_ = 0.5, r_0_ = 0.18, ρ = 0.1 and r_asympt_ = 1.05).

*Daphnandra repandula* (Laurales) has 16 tepals, eight stamens, seven staminodes and 11 carpels. The first two tepals are opposite. The other floral organs show an average divergence angle of 137.6° (±8.5°) suggesting that they are arranged in a Fibonacci spiral pattern. The perianth and androecium form pseudo-whorls of eight organs (Staedler and Endress 2009).

For both species convergence toward the attractor (i.e. the golden angle) is slower than for Fibonacci spirals and is cyclical (Fig. 22C and D).

### 4.7 Lucas phyllotaxis

The first few numbers in the Lucas sequence are 2, 1, 3, 4, 7, 11, …. Like in the Fibonacci sequence, each number is the sum of the two previous numbers, but the initialization is different. The evolution of Lucas phyllotaxis with the decrease in plastochron ratios over time is as follows: (1, 3) −> (3, 4) −> (4, 7) −> (7, 11) (Fig. 15). The divergence angle between successive primordia in meristems where the numbers of parastichies are terms of the Lucas sequence is close to the Lucas angle, which is equal to:

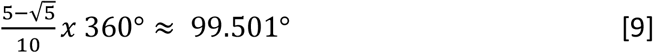

Marzec and Kappraff (1983) mathematically studied the effect of divergence angles on leaf distribution. They showed that certain remarkable angles optimize this distribution, meaning that they produce the densest meristems: they can all be expressed as a function of the golden ratio:

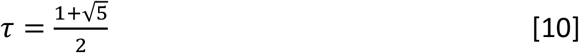

These angles are therefore all irrational, and we find among them the golden angle (137.507°) as well as the limit of the Lucas angle (99.501°):

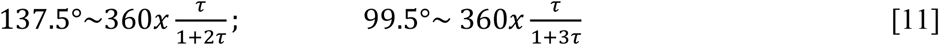

Strauss et al. (2020) showed that the golden angle (137.5°) optimizes light capture but other angles, such as the Lucas angle (99.5°) do so as well, which would go against the idea that the golden angle is favored by natural selection. They noted the similarity between the effects of the divergence angle on light capture and packing efficiency in a regular lattice, as calculated by Marzec and Kappraff (1983).

Church (p. 196 in Church 1904) lists *Sedum* (Crassulaceae), *Euphorbia* (Euphorbiaceae) and *Cereus* (Cactaceae) as examples of the (3, 4) mode (Fig. 76 and 77 p. 198 in Church 1904) and *Araucaria* (Araucariaceae) as an example of mode (7, 11), mode (4, 7) being very rare.

Rutishauser (1998) drew the cross section of *Sedum sexangulare*, which has the particularity of having 18 primordia in a (3, 4) phyllotaxis. The divergence angle is c. 101° and the plastochron ratio is 1.12 (both are average values for the first 18 primordia). *S. sexangulare* got its name because most members of this species have six orthostichies; however, some have not six but seven orthostichies (Deschatres 1954). This is the case for the specimen examined by Rutishauser (Fig. 23).

**Figure 23.**
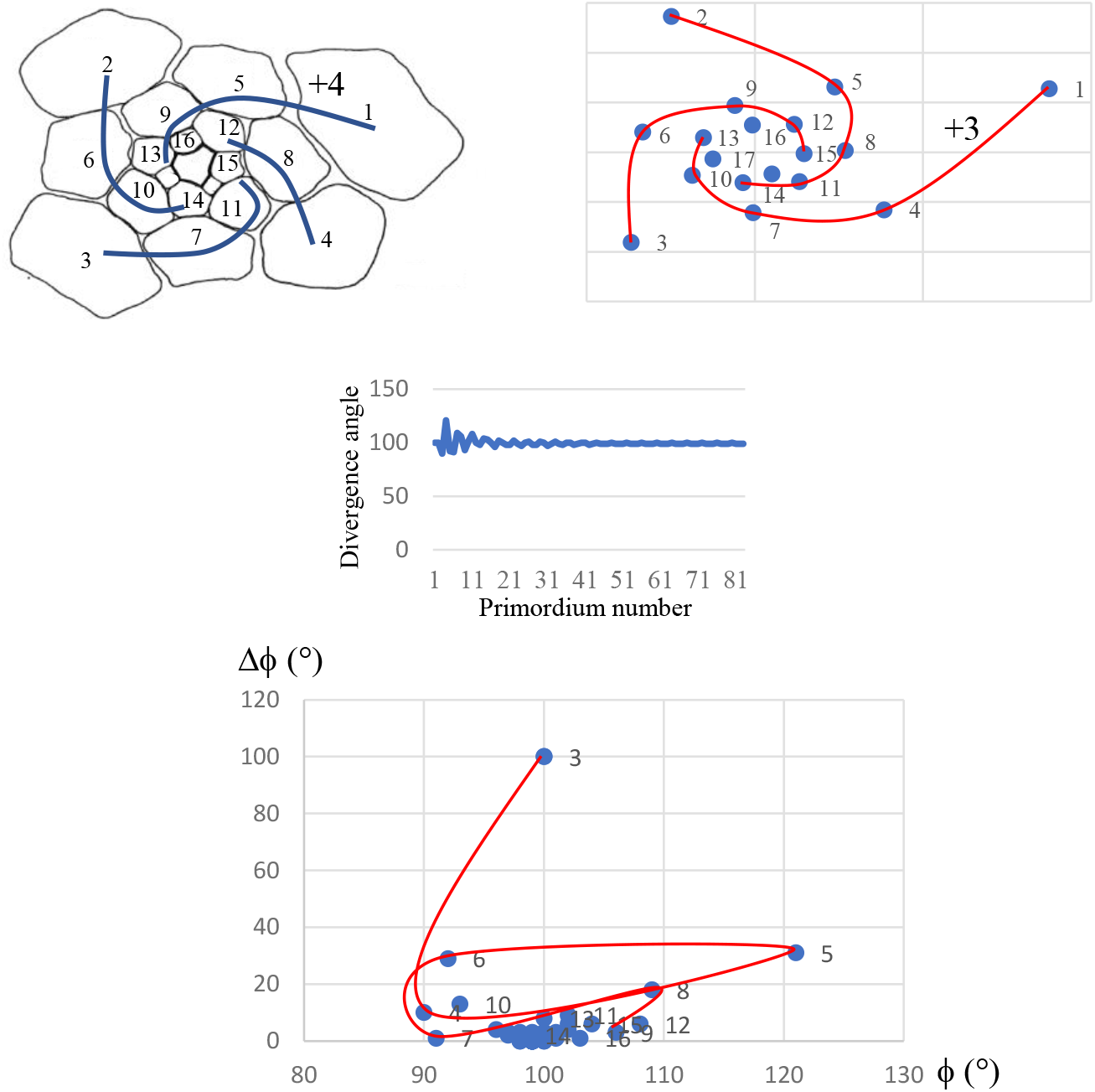
(A). The leaf apical meristem of the specimen of *Sedum sexangulare* examined by Rutishauser (1998). The phyllotaxis mode is (3, 4). Only the four clockwise parastichies are shown. (B) Centric representation of organ arrangement. (C) Divergence angle converging toward the Lucas angle. (D) Phase portrait of Lucas phyllotaxis. Parameter values are r_0_ = 0.4, ρ = 0.18 and r_asympt_ = 1.05.

Lucas phyllotaxis being as effective as Fibonacci phyllotaxis, one can wonder why the second occurs much more frequently in nature. According to Rutishauser (2019), this predominance is not due to natural selection, but to developmental constraints (‘laws of growth’) that force most spiral patterns to approach the famous Fibonacci angle. Indeed, the dynamics of phyllotaxic systems must be taken into account.

The transition from cotyledons to Lucas phyllotaxis, or any phyllotaxis mode other than Fibonacci phyllotaxis, requires crossing a potential barrier to reach a potential valley (see Fig. 15). Very specific initial conditions are necessary: for example, to model the leaf apical meristem of *S. sexangulare*, the divergence angle between each of the first three organs was set at 100° (Fig. 23). Furthermore, the size of the basin of attraction of Lucas phyllotaxis is very small. From their 1996 model, Douady and Couder also obtained (3, 4) and (4, 5) modes for specific values of the parameters.

Deschatres (1954) studied in detail the transition from a Fibonacci phyllotaxis ((2, 3) mode) to a Lucas phyllotaxis ((3, 4) mode) in *Sedum maximum* and used this as the basis for his criticism of the different “classical theories” based on divergence angles (this seems to change direction), Hofmeister’s rule (1868), or the first available space (Snow and Snow, 1948). From Deschatres point of view only Plantefol’s theory of multiple leaf helices (1949) would account for *Sedum maximum* specificity. However, none of the theories criticized by Plantefol still prevail.

Our static model (chapter 3 in Walch and Blaise 2023) applied to the stem of *Sedum maximum* faithfully reproduced the drawings made by Deschatres (Fig. 24).

**Figure 24.**
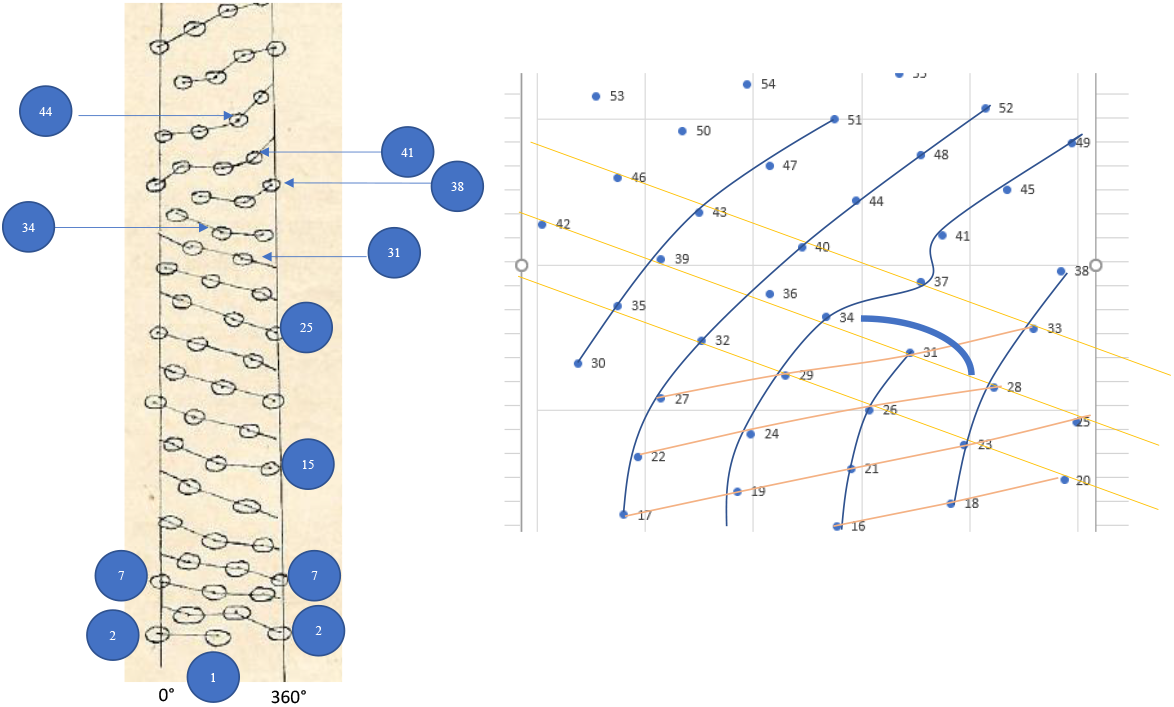
(A) Drawing of the stem of *Sedum maximum* by Deschartes (1954), with primordia numbered according to their initiation sequence. (B) Static model of the stem of *S. maximum.*

The static model allowed us to accurately calculate the average divergence angle between each leaf: around 138° at the base of the stem and 107° at the top. Pseudo-orthostichies of mode 2/5 (five leaves on a pseudo-orthostichy in two turns) were observed between leaves 6 and 26 and of mode 2/7 (seven leaves in two turns; ϕ ≈ 2/7 × 360° ≈ 103°) between leaves 38 and 52. Deschatres affirmed that: “the essential fact is the progressive transition to a disposition of the 2/7 series accompanied by an inversion of the generative spiral”.

According to Plantefol’s theory, the “leaf-generating centers” are arranged in helices “working in synchronism” in opposite or whorled leaves and “with an offset” in alternate leaves. The transition from a 2/5 series to a 2/7 series would highlight three progressively shifted leaf helices.

Careful observation of Deschatres’ drawings shows that one of the pseudo-orthostichies is interrupted at leaf 31 (Fig. 24B). The transition from mode (2, 3) to mode (3, 4) is therefore abrupt and not gradual. Such transitions have been observed by Wiss and Zagórska-Marek (2012) in *Magnolia*: a change in phyllotaxis pattern can occur with the disappearance or appearance of a parastichy, such as (8, 11) −> (7, 11) or (8, 12) −> (8, 13).

Zagórska-Marek (1987) attributes these transitions to discontinuities in the diameter of the apical meristem. The same type of dislocation is found in crystalline structures (edge dislocation) (Fig. 25).

**Figure 25.**
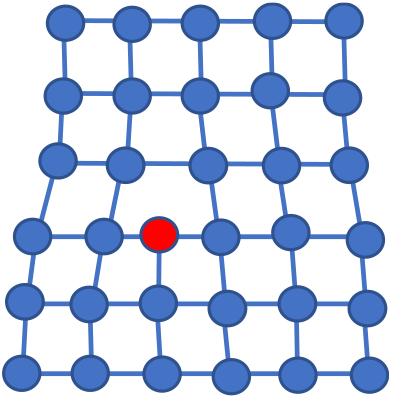
Edge dislocation in a crystalline material.

Here, the transformation is the consequence of a discontinuity in the sequence of plastochron ratios. Where dislocation takes place, the diameter of the meristem has not had time to adapt to the decrease in the number of organs on the front, so the d/D ratio (diameter of primordia over diameter of meristem) increases suddenly, as does the plastochron ratio.

## 5 A stochastic model

A stochastic model to account for biological noise in plant molecular processes was developed by Refahi et al. (2016). Noise can affect the delay between the initiation of two consecutive primordia (plastochron). By integrating stochasticity in the patterning system, their model produced occasional disrupted patterns that were similar to those seen in real plants. These errors can reveal information about how the signals that control phyllotaxis might work.

Here, we developed a stochastic model in which the plastochron ratio at the initiation of a new primordium is affected by a random process of amplitude ε. If this amplitude is 0.1, the plastochron ratios between two consecutive primordia can be randomly increased or decreased by a value between 0 and 0.1.

For average plasrochron ratios (Fig. 26A), spirals are not fundamentally modified although some primordia are permutated as evidenced by the periodic oscillations of the divergence angles between 137°, 223° (= 360°-137°) and 274° (= 2×137°). Such permutations were also found in the models of Refahi et al. (2016). This type of permutation has been observed in sunflower (Couder, 1998), *Arabidopsis thaliana* (Besnard et al., 2014), several other Brassicaceae showing spiral phyllotaxis as well as in monocot and dicot species from families such as Asparagaceae, Sapindaceae and Araliaceae (Refahi et al., 2016).

**Figure 26.**
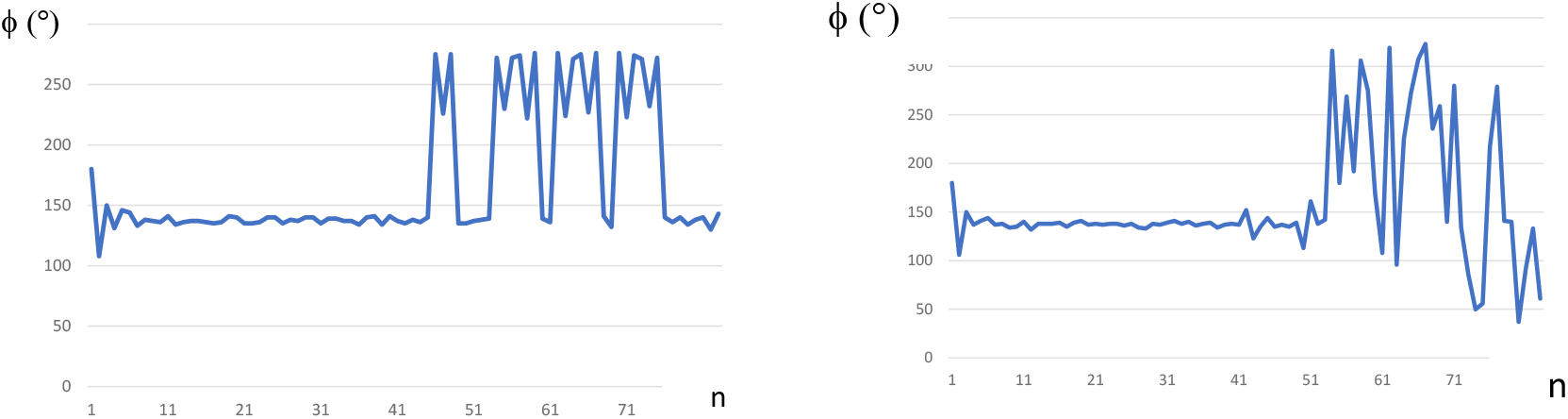
Divergence angle as a function of primordium number for r_0_ = 0.35; ρ = 0.18; ε= 0.12. (A) r_asympt_ = 1.05. (B) r_asympt_ = 1.02

When plastochron ratios are lower (r tending toward 1.02) (Fig. 26B), phyllotaxis becomes chaotic. Indeed, the amplitude of the random process being constant in this model over the entire course of development of the meristem, its effect will be greater when plastochron ratios are smaller.

## 5 Discussion

One of the major puzzles of phyllotaxis is that Fibonacci spirals are observed at a much higher frequency than other spirals such as Lucas spirals. In angiosperms, the Fibonacci series predominates (93.97% from a total of 8,720 observations) whereas the Lucas series is rare (only 1.18%) (Yin and Meicenheimer 2017).

Calculations by Marzec and Kapraff (1983) suggest that phyllotaxes that converge toward any noble (irrational) angle have equal packing performance. The question then arises of Darwinian selection seeming to favor Fibonacci phyllotaxis: the answer seems to be in the existence of developmental constraints independent of genetic or environmental variations that would limit organ positioning (Reinhart and Gola 2022; Godin et al. 2020).

Levitov’s model (1991) describes a structure composed of elements that repulse each other which is subjected to external mechanical stress. The purely geometric approach of van Iterson (van Iterson 1907; Godin et al. 2020) describes incompressible disks enclosed in a rigid container. The elements of Nisoli’s magnetic cactus (2009) are constrained to move only on a vertical axis. In Levitov’s model, the rise in the Fibonacci sequence is the consequence of an increase in the external pressure. For van Iterson, it is the consequence of a reduction in the radius of the disks compared to that of the container.

We have shown in a very general way that phyllotaxic spirals constitute a form of symmetry in the same way as axial symmetry (zygomorphy in botany). This symmetry is even more general than the one identified by Levitov, who assumes “strong compression” or any other form of stress of external origin. Thus, all arrangements of spiral symmetry belong to the same universality class.

This is the reason why they are found in different physical phenomena, such as in Douady and Couder’s magnetic experiment (1996), certain forms of Rayleigh-Bénard convection (Rivier et al. 1984), flux lattices in layered superconductors (Levitov, 1991) and the magnetic cactus (Nisoli et al. 2009).

In physics, these structures have no “finality”, but in plant biology, phyllotaxic spirals (irrespective of type) optimize the packing of building blocks (scales, primordia) which is enough to explain that natural selection has largely favored them. They should be compared to whorled phyllotaxes which are largely suboptimal when it comes to packing but have other advantages (Walch and Blaise, 2023).

Some models can be eliminated from the list of candidates if they make assumptions that are incompatible with botanical reality. This is the case of the van Iterson model, which assumes that primordia are enclosed in a fixed enclosure and that these primordia behave like “hard” disks. This is also the case of the Levitov model, which makes the assumption of an external pressure on the meristem that would be the control parameter of phyllotaxis. Their success is simply due to the fact that they are part of the spiral universality class.

By contrast, the control parameter in our model is the plastochron ratio, which is a measurable quantity, as shown by Rutishauser who measured this parameter in 17 plants (Rutishauser 1998). The equations of the model (with an exponential repulsion that decreases with distance) take into account biochemical inhibition but also contact pressure between “soft” organs after they are initiated by biochemical mechanisms. This internal contact pressure within the meristem is still part of the same spiral universality class (Walch and Blaise, 2023).

We have applied the theory of dynamical systems, an important branch of mathematics, to phyllotaxis. This clearly distinguishes whorled phyllotaxes with periodic dynamics from spiral phyllotaxes that converge toward an attractor.

The decrease of the plastochron ratio in spiral phyllotaxes results in dynamics evolving from a single attractor to multiple attractors (permutations of organs) to chaotic patterns, although this can resemble an apparent order in the form of “complex whorls”. This evolution is analogous to the evolution of fluid flows from laminar to turbulent.

The pseudo-whorled phyllotaxes observed in some Magnoliales and Laurales show slower convergence toward their attractor than spiral phyllotaxes, as well as damped cycles affecting the internal pseudo-whorls (i.e. stamens).

In principle, the developmental constraints specific to the biochemical inhibition mechanisms that control primordia initiation exclude any spiral phyllotaxis other than Fibonacci spirals. This explains the low frequency of Lucas phyllotaxes (and that of other noble angles), which, despite having the same selective advantage of Fibonacci spirals, should not even exist.

However, discontinuities in the plastochron ratios can allow plants to cross the potential barriers between Fibonacci phyllotaxis and other spiral phyllotaxes. In the case of *Sedum maximum*, the dislocation of a pseudo-orthostichy allows this barrier to be crossed and phyllotaxis to pass from a (3, 5) to a (3, 4) mode (i.e. a Lucas phyllotaxis mode), both of which optimizing packing.

According to Rutishauser (2019): « As already admitted by Darwin phyllotaxis patterns as observable in vascular plants appear as developmental patterns that are not under the control of natural selection. There are developmental constraints (‘laws of growth’) that force most spiral patterns to approach the famous Fibonacci angle, which is about 137.5°. »

Our conclusion is significantly different: natural selection favors all “noble” phyllotaxes, but developmental constraints almost exclude those that do not converge toward the golden angle. We thus agree with Strauss and al. (2020) who state that “the interplay of developmental constraints and potential fitness for the emergence of developmental patterns or complex traits” must be considered.

## Appendix

### Appendix 1

Let R_0_ be the radius of the central zone, V the centrifugal radial velocity and T the delay between the appearance of two primordia, then VT is the distance covered by a primordium during one plastochron. The growth index G = VT/R_0_ is a dimensionless number that characterizes the growth of the meristem irrespective of size and time scale.

If growth is exponential then the distance from the center of a primordium is:

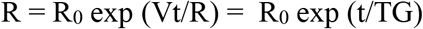

hence:

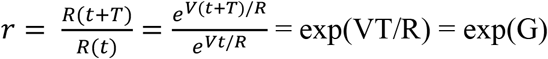

and the growth index G = ln(r).

### Appendix 2

The point K of coordinates (θk, hk) on a cylindrical lattice can be expressed by the complex number:

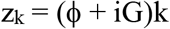

where k is the node number.

Applying the conformal transformation:

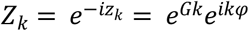

we obtain the algebraic form of the complex number Z_k_:

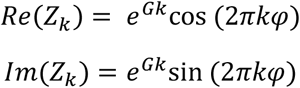

Primordia are located on a logarithmic spiral with the parametric equation:

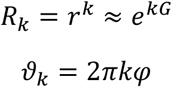

We eliminate k to obtain the polar equation of the spiral:

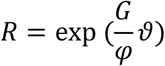

This logarithmic spiral is invariant for any transformation (similarity) consisting of a rotation of center O and angle a and a scaling of factor:

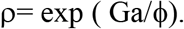

By this conformal transformation, a translation within the cylindrical lattice:

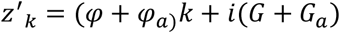

is transformed into a similarity within the centric network:

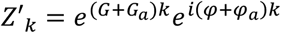

